# Peripheral CB1 receptor blockade acts as a memory enhancer through an adrenergic-dependent mechanism

**DOI:** 10.1101/2021.06.16.448227

**Authors:** Sara Martínez-Torres, Araceli Bergadà-Martínez, Jorge E. Ortega, Lorena Galera-López, Arnau Hervera, Antonio Ortega-Álvaro, Floortje Remmers, Emma Muñoz-Moreno, Guadalupe Soria, José Antonio del Río, Beat Lutz, Jose Ángel Ruíz-Ortega, J. Javier Meana, Rafael Maldonado, Andrés Ozaita

## Abstract

Peripheral inputs to the brain continuously shape its function and can influence the formation of non-emotional memory, but the underlying mechanisms have not been fully understood. Cannabinoid type-1 receptors (CB1R), widely distributed in the organism, is a well-recognized player in memory performance, and its systemic modulation significantly influences memory function. By assessing non-emotional memory in mice, we have now found a relevant role of peripheral CB1R in the formation of persistent memory. Indeed, peripherally restricted CB1R antagonism by using AM6545 showed a mnemonic effect that was occluded in adrenalectomized mice, after peripheral adrenergic blockade, or when vagus nerve was chemogenetically inhibited. Genetic CB1R deletion in dopamine β-hydroxylase-expressing cells enhanced the formation of persistent memory, supporting a role of peripheral CB1R modulating the adrenergic tone. Notably, brain connectivity was affected by peripheral CB1R inhibition, and *locus coeruleus* activity and extracellular hippocampal norepinephrine, were increased. In agreement, intra-hippocampal β-adrenergic blockade prevented AM6545 mnemonic effects. Together, we disclose a novel peripheral mechanism relevant for the modulation of the formation of persistent non-emotional memory.

## Introduction

Most everyday experienced events create non-emotional memories with different lifespans. They are generally rather short-lived and may fade away or may persist for longer periods, with the intervention of the hippocampus and further recall, even if all short- and long-lived memories were created from similar sensorial stimuli (Morris, 2006). Therefore, the persistence of stimuli-driven memories involves a discrimination/selection of worth-memorizing stimuli, which may occur around the time of encoding (Kandel et al., 2014; Yonelinas et al., 2019). Such memory persistence is enhanced in emotionally arousing experiences, where encoding is combined with the natural stress-coping response, which leads to a significant increase in memory persistence for the sensory information recorded at that time (De Quervain et al., 2016). In contrast, the mechanisms involved in memory persistence for non-emotional memories are not well understood, although they are also relevant for everyday life events.

The endocannabinoid system (ECS), highly expressed in the central nervous system (CNS) and peripheral tissues (Kano et al., 2009; Maccarrone et al., 2015), plays a key role in learning and memory (Kano et al., 2009). The cannabinoid type-1 receptor (CB1R) is strongly expressed in the brain (Pacher et al., 2006), predominantly localized at presynaptic sites of different neuronal cell types, where it suppresses neurotransmitter release depending on local synaptic activity (Castillo et al., 2012). Exogenous compounds with agonist properties for CB1R contribute to memory impairment (Niyuhire et al., 2007; Puighermanal et al., 2012) in mice in non-emotional memory medels, such as the novel object-recognition test (NORT), while pharmacological or genetic CB1R blockade increases memory persistence in these models (Maccarrone et al., 2002; Reibaud et al., 1999). Although the mechanisms involved are largely unknown, such regulation of memory by CB1R blockade was previously assumed to occur solely through centrally located receptors (Zanettini et al., 2011).

We set to challenge this dogma by studing the mnemonic effect of a peripherally restricted CB1R antagonist, AM6545 (Tam et al., 2010). Peripheral CB1R had been previously found relevant for the amnesic effects associated to stress at the time of memory consolidation (Busquets-Garcia et al., 2016), but their contribution to memory enhancement had not been previously assessed.

We found that peripheral blockade of CB1R on adrenergic cells enhanced non-emotional memory persistence through an adrenergic mechanism involving the vagus nerve. Under these conditions, brain connectivity was increased and hippocampal norepinephrine release was boosted as a plausible mechanism underlying this enhancement of memory persistence.

## Results

### CB1R inhibition enhances memory persistence in the novel object-recognition memory test

Novel object-recognition memory is a labile non-emotional type of memory usually evaluated 3 h or 24 h after the familiarization session, when mice readily discriminate novel and familiar objects (Fig. 1a). Notably, discrimination values significantly decrease when novel object-recognition memory is assessed 48 h after the familiarization phase (one-way ANOVA, interaction: F (2,19) = 8.55, p = 0.002; *post hoc* Tukey, 3 h vs 48 h p = 0.003; 24h vs 48h p = 0.007) (Fig. 1a). We used this non-emotional memory paradigm assayed 48 h after the familiarization phase to evaluate the role of CB1R inhibition in memory persistence. We found that mice with acute post-familiarization treatment with a low dose of the systemic CB1R selective antagonist rimonabant (1 mg/kg, i.p.) showed higher memory persistence than vehicle-treated mice (Student’s t-test: p = 0.02) (Fig. 1c). In addition, heterozygous mice for the *Cnr1* gene (CB1HZ) also showed enhanced memory persistence (Student’s t-test: p = 0.004) compared to their wild-type littermates (Fig. 1d), indicating that such a modulation in memory persistence is CB1R dependent. Post-familiarization administration of the peripherally-restricted CB1R antagonist AM6545 (1 mg/kg, i.p.) also enhanced novel object-recognition memory at 48 h (Student’s t-test: p = 0.002) (Fig. 1e) pointing to the involvement of a peripheral mechanism. No differences in total exploration time were detected between genotypes or pharmacological treatments in any of the experimental groups above (Supplementary Fig. 1) discarding a possible bias due to differences between groups in exploratory behavior. Furthermore, AM6545 treatment did not affect locomotor activity analyzed for 120 min post-administration (Supplementary Fig. 2) excluding a major unspecific effect of the treatment on behavioral responses.

**Fig. 1.**
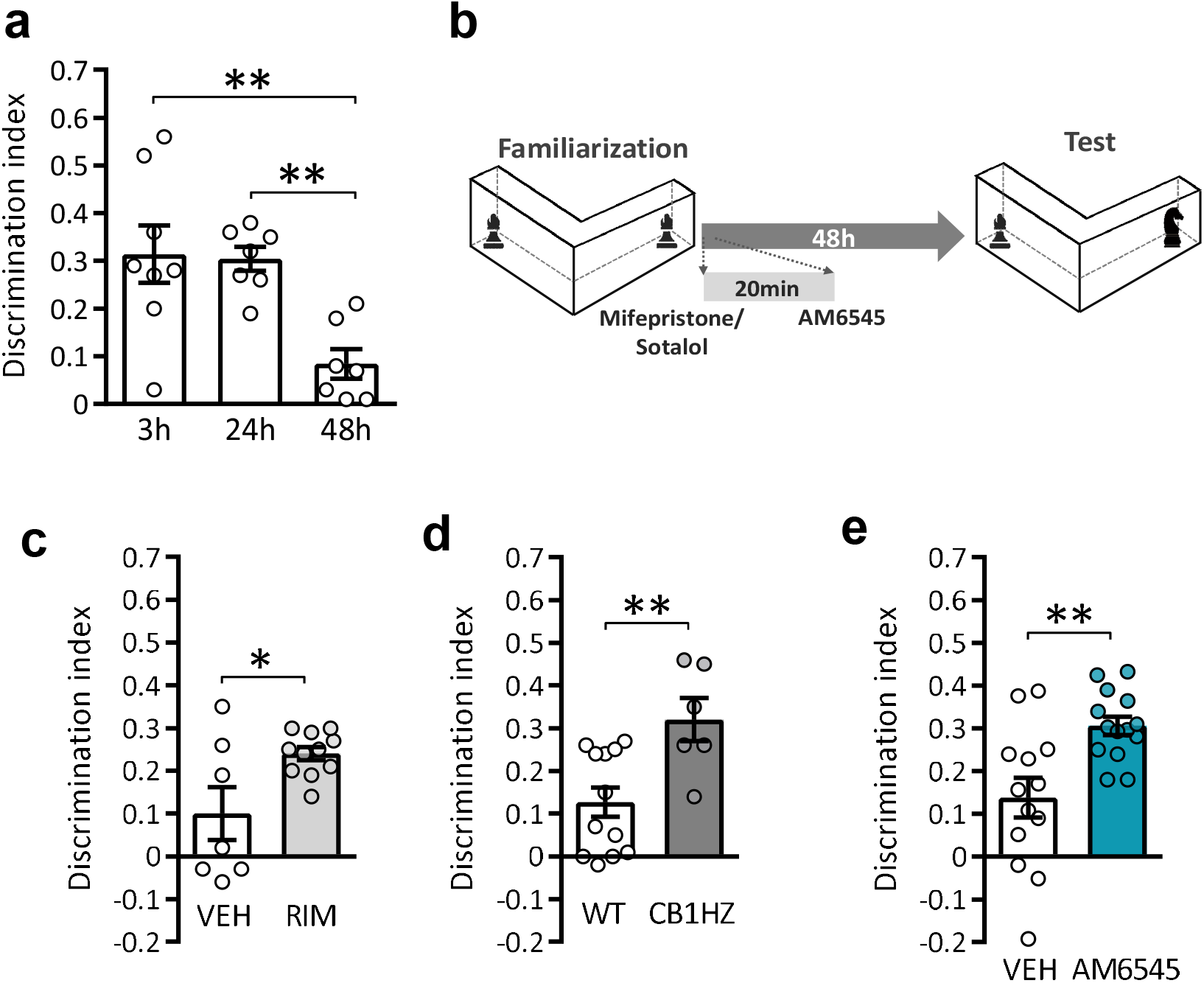
Pharmacological or genetic inhibition of CB1R improves memory persistence in the novel object-recognition test. **a** Discrimination index values obtained at 3 h, 24 h and 48 h after the familiarization phase (n = 5 - 8). **b** Schematic representation of pre-treatment and treatment after the familiarization phase. **c-e** Discrimination index values in NORT at 48 h **c** after acute post-familiarization treatment with vehicle (VEH) or rimonabant (RIM) (1 mg/kg) (n = 7 - 11), **d** in CB1HZ and WT mice (n = 6 - 8), **e** after acute post-familiarization treatment with vehicle (VEH) or AM6545 (1 mg/kg) (n = 7 - 8). Data are expressed as mean ± s.e.m. *p < 0.05, **p < 0.01 by one-way ANOVA test followed by Tukey *post hoc* or Student’s t-test.

### Enhanced memory persistence of peripheral CB1R inhibition involves a peripheral adrenergic mechanism

We hypothesized that a peripherally located tissue, such as the adrenal glands which express CB1R (Hillard, 2015), could be a relevant player modulating memory consolidation (McIntyre et al., 2012). Therefore, we evaluated the effect of post-familiarization AM6545 administration in bilaterally adrenalectomized mice. Memory persistence enhancement by AM6545 was significantly reduced in mice without adrenal glands (two-way ANOVA, interaction: F (1,22) = 4.89, p = 0.037; *post hoc* Tukey, naive-VEH vs naive-AM6545, p = 0.022; naive-AM6545 vs ADX-AM6545, p = 0.026) (Fig. 2a), supporting the role of CB1R blockade in this paticular peripheral tissue. Adrenal glands release glucocorticoids and catecholamines into the blood, both relevant for memory (McIntyre et al., 2012). In addition, both rimonabant (Wade et al., 2006) and high doses of AM6545 have been observed to increase circulating corticosteroids (Roberts & Hillard, 2020). To elucidate out which hormones are responsible for the mnemonic effects produced by peripheral CB1R blockade, mice were pre-treated after the familiarization phase with the glucocorticoid receptor antagonist mifepristone (50 mg/kg i.p.) or the peripherally-restricted β-adrenergic receptor antagonist sotalol (10 mg/kg i.p.) 20 min before AM6545 injection (Fig. 2b). Mifepristone pre-treatment did not prevent enhancement of memory persistence by AM6545 (two-way ANOVA, interaction: F (1,21) = 0.038, p = 0.845; mifepristone/vehicle effect: F (1,21) = 0.707, p = 0.409; AM6545/vehicle effect: F (1,21) = 25.11, p<0.001) (Fig. 2b). In contrast, mice pre-treated with sotalol did not show the memory improvement observed in AM6545-treated mice (two-way ANOVA, interaction: F (1,31) = 7.58, p = 0.009; *post hoc* Tukey, Saline-VEH vs Saline-AM6545 p = 0.01; Saline-AM6545 vs Sotalol-AM6545 p = 0.001) (Fig. 2c).

**Fig. 2.**
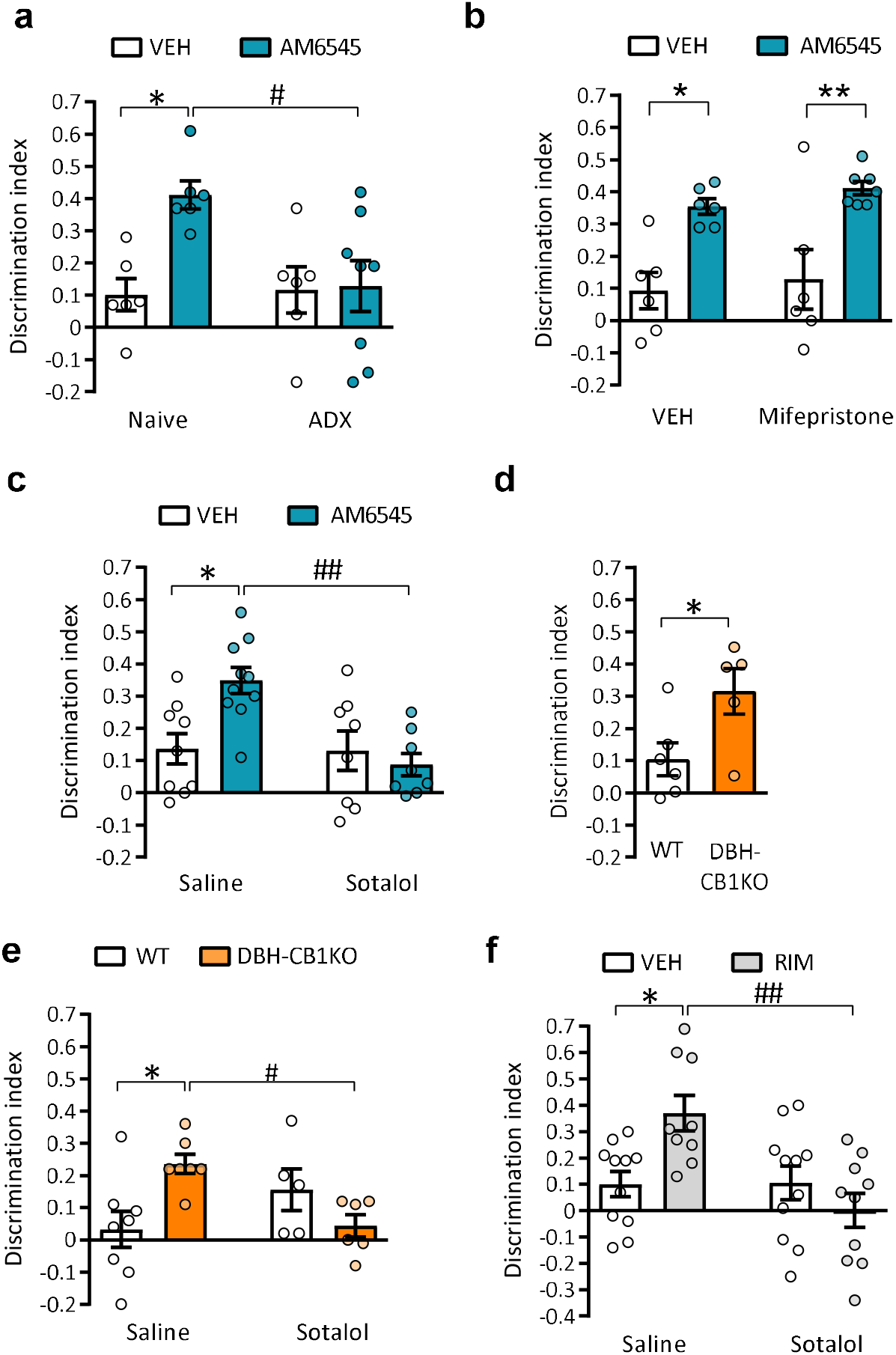
AM6545 enhances novel object-recognition memory through a peripheral β-adrenergic mechanism. **a** Discrimination index values obtained in the NORT performed at 48 h of adrenalectomized (ADX) or naive mice treated with vehicle (VEH) or AM6545 (1 mg/kg) (n = 6 - 8). **b, c** Discrimination index values obtained in the NORT performed at 48 h of mice treated with vehicle (VEH) or AM6545 (1 mg/kg) after pre-treatment with **b** vehicle (VEH) or mifepristone (50 mg/kg) (n = 6 - 7) and **c** saline or sotalol (10 mg/kg) (n = 8 - 10). **d** Discrimination index values in WT or DBH-CB1KO mice in NORT at 48 h (n = 5 - 6) **e** and after saline or sotalol (10 mg/kg) treatment (n = 6 - 8) **f** Discrimination index values for mice pre-treated with saline or sotalol (10 mg/kg) prior to rimonabant (RIM) (1 mg/kg) or vehicle (VEH) in the NORT at 48 h (n = 9 - 11). Data are expressed as mean ± s.e.m. *p < 0.05, **p < 0.01 (treatment effect) ^#^p < 0.05, ^##^p < 0.01 (pre-treatment effect) by two-way ANOVA test followed by Tukey *post hoc*.

In the light of these data, we assessed whether inhibition of CB1R exclusively in dopamine β-hydroxylase cells (DBH+ cells), the main cells responsible for circulating levels of epinephrine/norepinephrine, could mimic the mnemonic effect of systemic and peripheral CB1R antagonists. We used a combination of genetic and pharmacological approaches to first show that conditional knock-out mice lacking the CB1R in DBH+ cells (DBH-CB1KO mice) displayed enhanced novel object-recognition memory persistence compared to wild-type controls (Student’s t-test: p = 0.04) (Fig. 2d). In addition, enhanced memory persistence in DBH-CB1KO mice was abolished by sotalol administration (two-way ANOVA, interaction: F (1,22) = 10.47, p = 0.003; *post hoc* Tukey, Saline-WT vs Saline-DBH-CB1KO p = 0.01; Saline-DBH-CB1KO vs Sotalol-DBH-CB1KO p = 0.04 (Fig. 2e), pointing to a relevant role of CB1R-modulated peripheral adrenergic/noradrenergic tone in memory persistence. Sotalol pre-treatment similarly prevented the cognitive improvement elicited by systemically-acting rimonabant supporting the relative prominence of peripheral CB1R blockade in this effect of rimonabant (two-way ANOVA, interaction: F (1,37) = 9.408, p = 0.004; *post hoc* Tukey, Saline-VEH vs Saline-rimonabant p = 0.01; Saline-rimonabant vs Sotalol-rimonabant p = 0.001) (Fig. 2f). No differences in total exploration time were detected in the memory test between genotypes or pharmacological treatments in any of the experimental groups above (Supplementary Fig. 3).

### Chemogenetic vagus nerve inhibition prevents memory improvement produced by peripheral CB1R blockade

Systemic epinephrine administration does not cross the blood brain barrier and enhances hippocampal dependent memory in rodents by the activation of peripheral β-adrenergic receptors (Dornelles et al., 2007). Such mnemonic effects have been hypothesized to occur either by the activation of β-adrenergic receptors in the liver and the subsequent increase of blood glucose levels, or by the activation of β-adrenergic receptors on the afferent fibers of the vagus nerve (Gold & Korol, 2012).

To further elucidate whether either of these two mechanisms could be involved in the mnemonic effects produced by acute AM6545 administration, we first measured blood glucose levels after AM6545 administration in mice. No differences in blood glucose levels were observed in mice treated with AM6545 in comparison to vehicle-treated group (Fig. 3a).

**Fig. 3.**
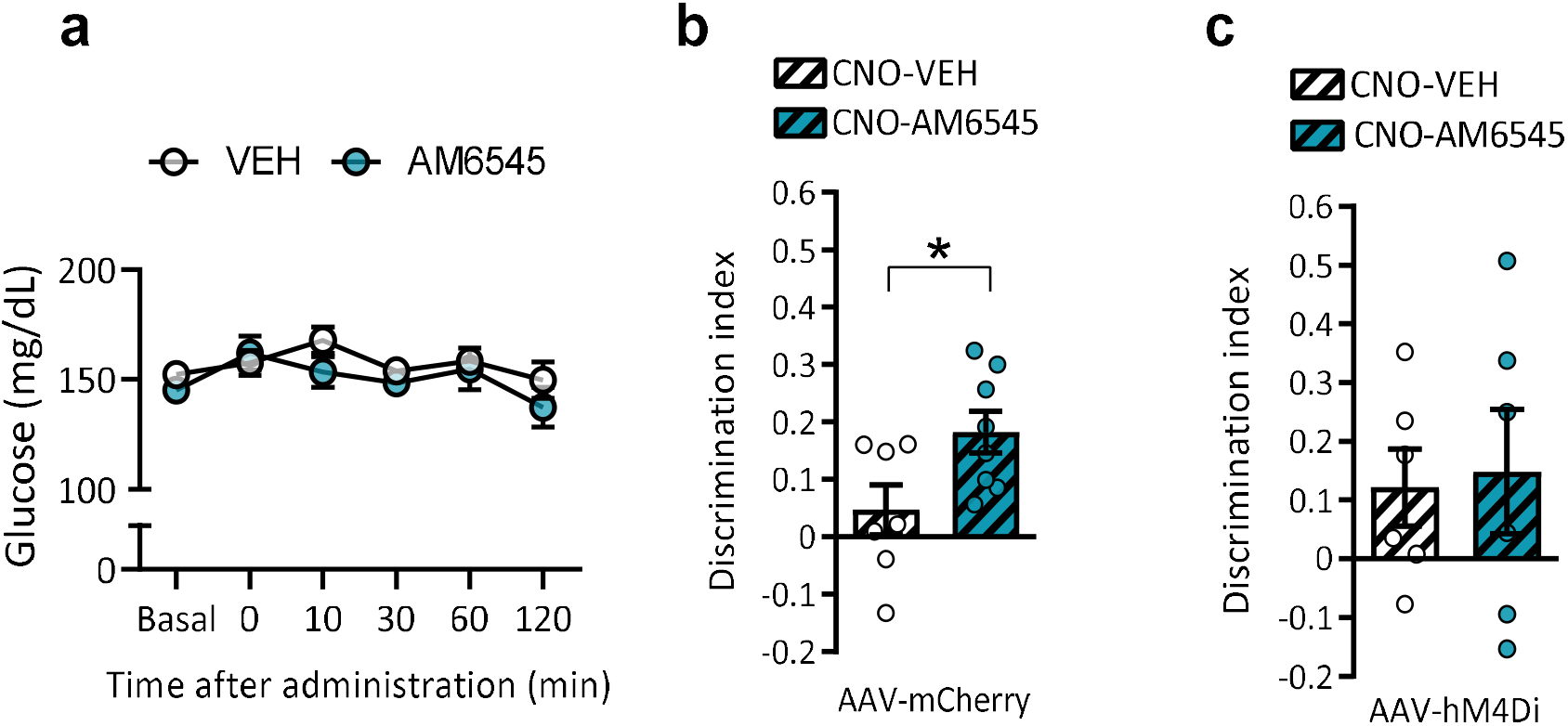
AM6545 memory improvement depends on vagus nerve activation. **a** Glucose blood levels for 120 min after acute vehicle (VEH) or AM6545 (1 mg/kg) administration in mice (n = 6 – 7) **b**,**c** Discrimination index values in NORT 48 h after acute vehicle (VEH) or AM6545 (1 mg/kg) and CNO (3 mg/kg) administration in mice infected with **b** AAV5-mCherry-control or **c** AAV5-mCherry-hM4Di in the vagus nerve (n = 6 – 7). Data are expressed as mean ± s.e.m. *p < 0.05 by Student’s t test or two-way repeated measures ANOVA test.

We used a chemogenetic approach to selectively reduce the neural activity of vagus nerve and assess the involvement of vagal fibers as a major link between peripheral AM6545 effects and memory performance We first confirmed that AM6545 effect on memory persistence was maintained in control mice receiving bilateral vagus nerve injection of AAV5-mCherry and subsequently treated with clozapine N-oxide (CNO, 3 mg/kg, i.p.) (Fig. 3b), indicating that surgery, vagus nerve infection with viral vectors, and CNO administration were compatible with the assessment of memory persistence. However, chemogenetic inhibition in animals with bilateral vagus nerve injection of AAV5-hM4Di prevented AM6545-memory enhancement (Fig. 3c) revealing a role of vagus nerve afferents in the mnemonic effect of AM6545. As expected, no differences were observed neither when memory was evaluated 24 h after familiarization in AAV5-hM4Di mice nor in exploratory behavior during the memory test (Supplementary Fig. 4). Thus, AM6545 administration improves memory consolidation through an adrenergic mechanism involving the vagus nerve.

### Central effects of AM6545 administration

We first analyzed the pattern of c-Fos expression, as a marker of neuronal activity, focusing on brain regions relevant for memory performance. Samples were obtained 90 min after receiving AM6545 or vehicle once mice has finished the familiarization phase in the NORT. Brain areas analysed included CA1 and CA3 hippocampal regions, dentate gyrus, prelimbic and infralimbic prefrontal cortex, locus coeruleus and basal, lateral and central regions of the amygdala. No significant differences in c-Fos expression were observed between AM6545- and vehicle-treated mice in the areas analyzed (Supplementary Fig.5). However, network analysis using the Pearson correlation coefficient for each pair of regions revealed that AM6545 treatment produced a significant alteration in the functional connectivity of these brain areas compared to the vehicle condition. (Fig. 4a-b and Supplementary Fig.6). We calculated Z-scores starting from the Pearson *r* correlation values to compare positive and negative connectivity between groups. AM6545 treatment did not significantly modified neither positive nor negative correlations (Fig. 4c-d).

**Fig. 4.**
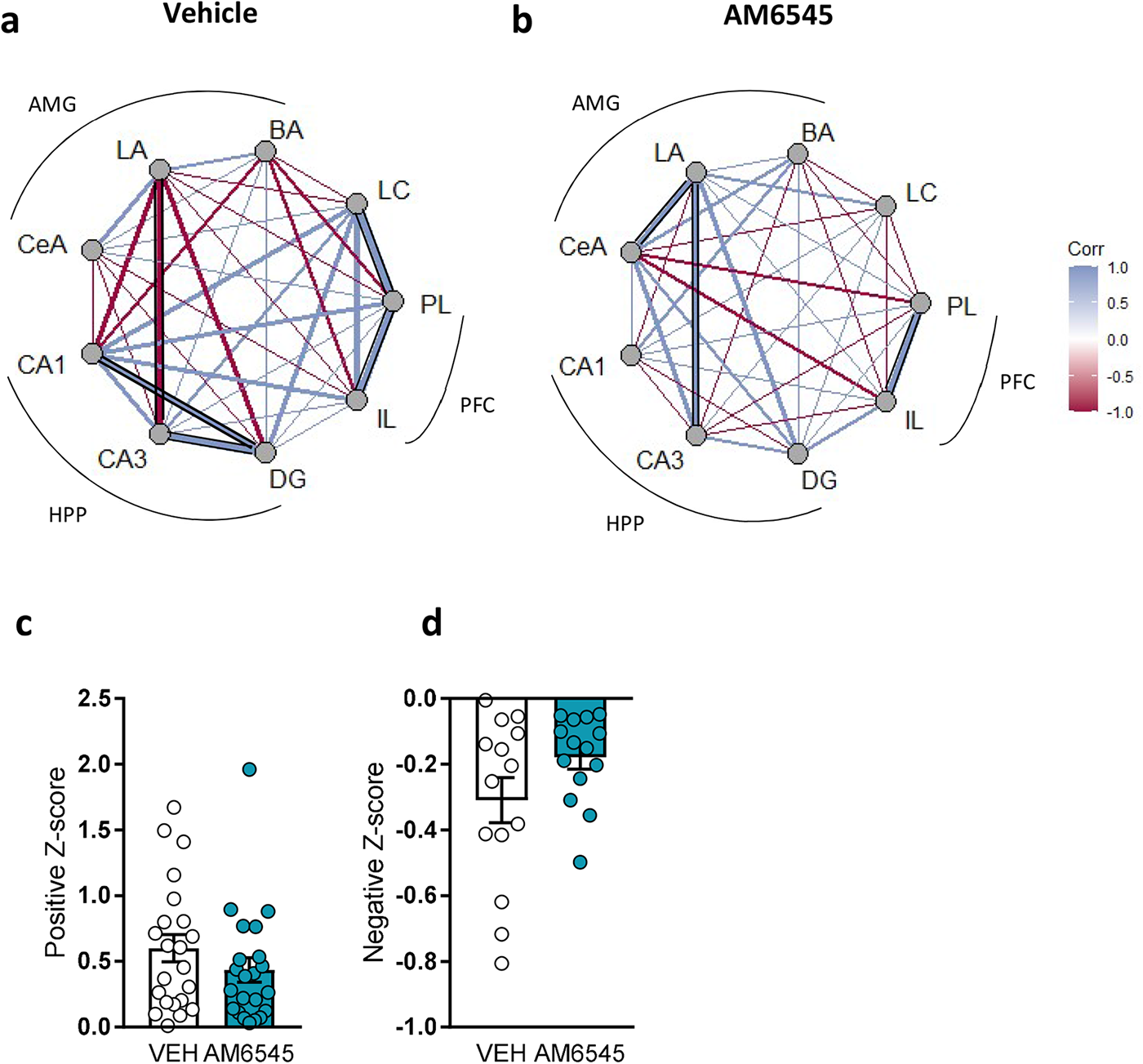
Circos plot and z-score analysis of c-Fos expression correlation analysis in different brain regions after acute AM6545 administration. Network graphs of c-Fos expression based on the Pearson coefficient in **a** vehicle and **b** AM6545 (1 mg/kg) between different brain areas, including prelimbic cortex (PL), infralimbic cortex (IL), *cornu ammonis* 1 (CA1), dentate gyrus (DG), *cornu ammonis* 3 (CA3), *locus coeruleus* (LC), basal amygdala (BA), central amygdala (CeA) and lateral amygdala (LA). Significant correlations are depicted with black lines. **c**,**d** Z-score values of the **c** positive and **d** negative Pearson *r* correlation coefficients between the different brain regions analysed.

We further explored the central effects after peripheral CB1R blockade by performing resting state functional magnetic resonance imaging (rsfMRI). After 1 hour of AM6545 administration, global connectivity metrics showed a significant enhancement of AM6545 treatment in whole brain local efficiency (Fig. 5a) (uncorrected p<0.05) indicating an enhanced ability of parallel transmission of information. Regional connectivity analysis showed significant increases of nodal efficiency values in AM6545-treated mice compared to vehicle-treated mice in left frontal cortex (Kruskal-Wallis: p<0.01), right hippocampus (Kruskal-Wallis: p = 0.023) and right globus pallidus (Kruskal-Wallis: p = 0.023) (Fig. 5 b-d), although these differences did not reach significance when corrected for multiple comparisons (FDR), indicating a subtle effect of AM6545 enhancing brain connectivity. Seed-based analysis of rsfMRI with seed in the posterior brainstem, which contains the nucleus tractus solitarius (NTS), a region receiving vagal afferents, exhibited a different pattern of connectivity in AM6545-treated animals compared to vehicle-treated animals. Significant differences of this connectivity pattern were observed in areas corresponding to hippocampus, frontal cortex, cingulate cortex, brainstem nuclei such as pontine reticular nucleus, several preolivary and spinal trigeminal nuclei and cerebellar nuclei (Fig. 5e). In these regions, AM6545-treated animals showed strong negative correlation values with the seed BOLD signal, especially in hippocampus, that are not observed in vehicle treated animals (Supplementary Fig.7).

**Fig. 5.**
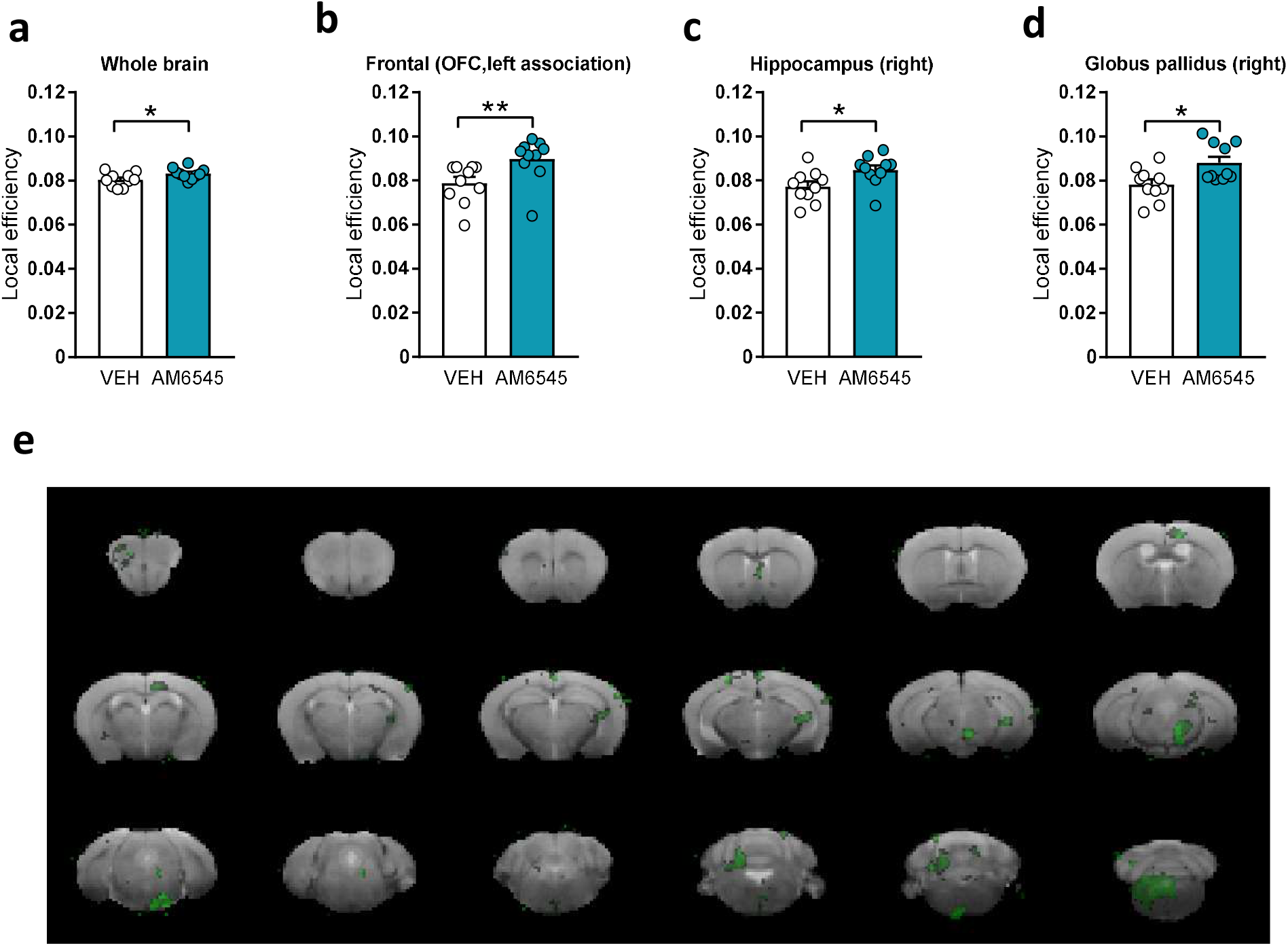
Effects of AM6545 administration on brain functional connectivity by resting state functional magnetic resonance imaging (rsfMRI). **a** Whole brain local efficiency and **b-d** nodal efficiencies of **b** left frontal cortex, **c** right hippocampus and **d** right globus pallidus. **e** Statistical map showing significant differences between connectivity of posterior brainstem with the rest of the brain in vehicle and AM6545 treated animals (comparison vehicle>AM6545; p<0.01). Data are expressed as mean ± s.e.m. *p < 0.05, **p < 0.01 by Kruskal-Wallis test.

### Increased hippocampal norepinephrine mobilization by peripheral CB1R blockade induces memory improvement

Next, we measured the firing rate at *locus coeruleus* neurons after CB1R peripheral modulation with AM6545. Systemic AM6545 produced a non-significant enhancement of basal firing rates compared to vehicle (Fig. 6a). To assess the functional significance of the basal firing rate, we performed extracellular microdialysis analysis in the hippocampus after systemic AM6545 treatment. Analysis of norepinephrine (NE), dopamine (DA) and serotonin (5-HT) extracellular levels after AM6545 administration revealed a specific transient increase in NE in comparison to the vehicle-treated mice (two-way repeated measures ANOVA, interaction: F (1,14) = 2.19 p = 0.009) (Fig. 6b and Supplementary Fig.8). We then tested whether β-adrenergic receptors in the hippocampal region were involved in the increased memory persistence mediated by AM6545. We found that local intra-hippocampal microinjection of propranolol at a dose that did not affect memory performance on its own (1 µg per 0.5 µL per side, see Supplementary Fig. 9) blocked the mnemonic effects produced by systemic AM6545 administration (two-way ANOVA, interaction: F (1,119) = 5.03, p = 0.03; *post hoc* Tukey, Saline-VEH vs Saline-AM6545 p = 0.004; Saline-AM6545 vs Propranolol-AM6545 p = 0.01) (Fig. 6c), without affecting the exploratory behavior on the memory test (Fig. 6d). These data indicate the functional relevance of noradrenergic hippocampal activation in the effect of peripheral CB1R blockade on memory persistence.

**Fig. 6.**
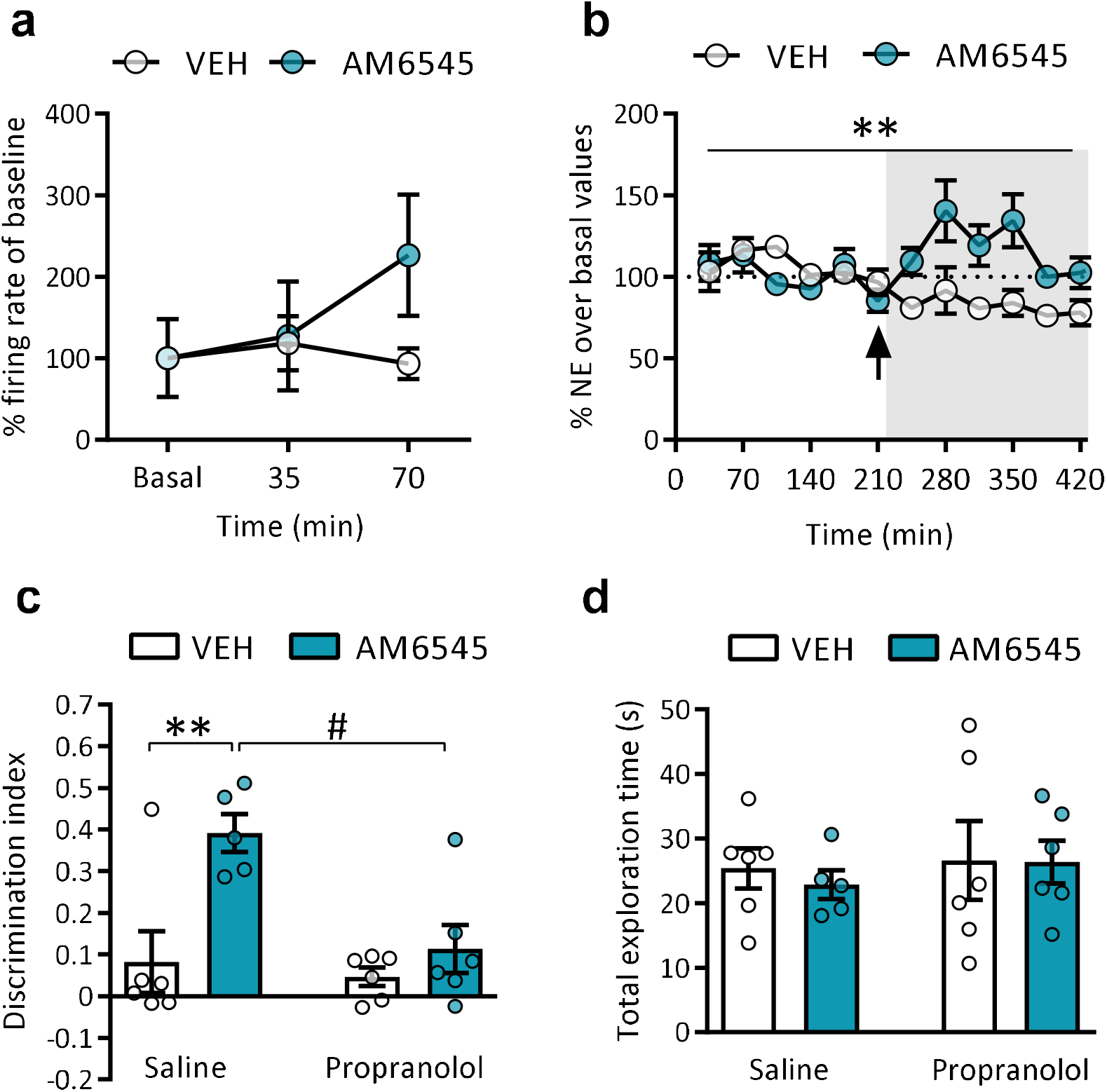
Acute AM6545 treatment increases central noradrenergic activity. **a** Percentage of mean firing rate in the LC after acute vehicle (VEH) or AM6545 (1 mg/kg) administration respect to baseline values (n = 2 - 3) **b** Percentage of extracellular norepinephrine (NE) levels in the hippocampus after acute vehicle (VEH) or AM6545 (1 mg/kg) administration relative to baseline values (n = 6 - 7). The arrow indicates the time of administration. **c** Discrimination index values and **d** total exploration time obtained in the NORT performed at 48 h mice treated with AM6545 (1 mg/kg) or vehicle (VEH) after bilateral intrahippocampal injection of saline or propranolol (1 μg/μl 0.5 μL per side) (n = 5 - 6). Data are expressed as mean ± s.e.m. For microdialysis **p < 0.01 two-way repeated measures ANOVA test followed by Tukey *post hoc* was performed. For NORT **p < 0.01 (treatment effect) ^#^p < 0.05 (pre-treatment effect) by two-way ANOVA test followed by Tukey *post hoc*. For the firing rate at *locus coeruleus* statistical significance was calculated by two-way repeated measures ANOVA test.

## Discussion

Our study identifies a relevant role of the peripheral ECS in modulating non-emotional memory persistence through the mobilization of central and peripheral adrenergic/noradrenergic mechanisms. We chose to study novel object-recognition memory, a hippocampal-dependent test (Cohen & Stackman, 2015) since this is a model of non-emotional memory and memory persistence might be modulated in this test by post-training manipulation. In this regard, we have previously observed deficits in novel object-recognition memory by CB1R agonists (Puighermanal et al., 2009), endocannabinoid build-up (Busquets-Garcia et al., 2011) and stress (Busquets-Garcia et al., 2016), all interventions after the familiarization phase. In the present study, we analyzed novel object-recognition memory recall 48 h after the familiarization phase since at that time mice usually show reduced signs of novel object discrimination compared to shorter intervals. Using this 48 h interval, we found that overall CB1R blockade, through pharmacological or genetic approaches, significantly increased discrimination indexes. In agreement with this, a number of previous studies had also demonstrated that both pharmacological and genetic CB1R inactivation enhanced memory in different hippocampal-dependent tasks (Jacob et al., 2012; Lichtman, 2000; Maccarrone et al., 2002; Reibaud et al., 1999; Takahashi et al., 2005; Wolff & Leander, 2003). We found that the peripherally restricted CB1R neutral antagonist AM6545 administered after the familiarization phase significantly enhanced object-recognition memory persistence in mice. This drug does not show significant blood-brain barrier permeability at doses even ten times higher than the one used in the present study (Tam et al., 2010), suggesting that the AM6545 effects derive in our study from a peripheral mechanism. Notably, adrenalectomized mice treated with AM6545 did not present an enhancement in memory persistence, pointing to a relevant role of adrenal glands in this response. Epinephrine/norepinephrine and corticosteroids secreted by the adrenal glands have a significant impact in memory consolidation (McIntyre et al., 2012; Roozendaal & McGaugh, 2011; Yang et al., 2013). We then gathered evidence that pointed to the mobilization of the adrenergic transmission by peripheral CB1R blockade, but not to the mobilization of corticosteroids, given that AM6545-induced memory persistence enhancement was prevented by the peripherally restricted β-adrenergic receptor antagonist sotalol, but not by the corticosteroid receptor antagonist mifepristone. Such differential modulation by adrenal hormones of AM6545 effect is indicative of a rather specific mechanism of peripheral CB1R blockade for potentiating the consolidation of non-emotional memory. Indeed, sotalol has been previously found ineffective in modulating fear conditioning (Lee et al., 2001) and mifepristone was found to reduce contextual memory in a fear conditioning paradigm (Zhou et al., 2010), whereas it did not modify NORT results in our experimental conditions. Notably, mice lacking CB1R in DBH+ cells (Busquets-Garcia et al., 2016) (DBH-CB1KO) reproduced a similar phenotype on enhanced memory persistence as constitutive CB1HZ mice. Our present finding that sotalol prevents such a mnemonic phenotype in DBH-CB1KO mice and after systemic rimonabant treatment, points to a relevant effect of the peripheral adrenergic CB1R in the systemic effect of CB1R inhibition, and a hitherto undisclosed role of peripheral CB1R signaling in memory persistence.

The study of c-Fos expression after AM6545 administration allowed to confirm no major differences in neuronal stimulation due to the treatment, but significant alterations in functional connectivity between areas analyzed. Although c-Fos may not reveal the real activity of brain regions given its limited expression and the time constraints of the analysis (McReynolds et al., 2018), this approach has been successfully performed to acquire a glimpse of brain functional connectivity between discrete areas (Silva et al., 2019; Tanimizu et al., 2017). The subtle alterations in brain connectivity revealed by our biased c-Fos inter-regional correlation analysis were further corroborated by rsfMRI using an unbiased approach to study connectivity. The brain’s intrinsic functional organization associated to the resting state has been proposed to determine the ability to create flexible and suitable behavioral outcomes to cognitive demands (Sala-Llonch et al., 2012). Indeed, enhancements in nodal efficiency, a representation for the ability of information propagation in the subnetwork of regions connected to a node, were observed specifically in the frontal cortex, hippocampus and globus pallidus, a result in agreement with the engagement of these regions in long-term memory consolidation (Kitamura et al., 2017; Tanimizu et al., 2017) promoted by AM6545 treatment.

A growing body of evidence has demonstrated the efficacy of systemic epinephrine administration, which does not cross the blood-brain barrier, to enhance hippocampal-dependent memory in rodents (Dornelles et al., 2007; Talley et al., 2000) by enhancing central noradrenergic signaling. Our data reveal that AM6545 administration did modestly modify brain activity to produce a reduction in brain network connectivity and a partial increase in basal LC firing and norepinephrine extracellular levels in the hippocampus, suggesting increased activity of LC projections to this brain region. These finding are compatible with the activation of the vagus nerve by increasing peripheral adrenaline/noradrenaline, since it was demonstrated that epinephrine administration results in increased vagal nerve firing that was sensitive to sotalol administration (Miyashita & Williams, 2006). Vagus nerve stimulation has been found to enhance memory retention in mouse models (Vázquez-Oliver et al., 2020) as well as in humans (Ghacibeh et al., 2006). LC activity, partially determined by the vagal nerve afferents, is well stablished to be physiologically engaged on novelty and to prime the persistence of hippocampal-based long-term memories (Hansen, 2017; Sara, 2009). Moreover, norepinephrine in the hippocampus facilitates the storage of new memories through the regulation of neural excitability and synaptic plasticity (Hagena et al., 2016). Although norepinephrine can bind to both α- and β-adrenergic receptors, synaptic information and plasticity in the hippocampus is proposed to depend largely on the activation of β-adrenergic receptors (Kemp & Manahan-Vaughan, 2008). Indeed, LC stimulation modulates hippocampal synaptic strength (Hansen & Manahan-Vaughan, 2015; Kemp & Manahan-Vaughan, 2012, 2008), spike coupling of CA1 pyramidal neurons (Bacon et al., 2020) and improves memory through a β-adrenergic-dependent mechanism (Hagena et al., 2016; Hansen & Manahan-Vaughan, 2015). In agreement, we observed that the intra-hippocampal blockade of β-adrenergic receptors with propranolol prevented the increase of novel object-recognition memory persistence produced by AM6545.

Altogether, our study identifies peripheral CB1R as a relevant target to enhance non-emotional memory persistence through a central and peripheral adrenergic mechanism that engages the vagus nerve. This novel peripheral mechanism involving CB1R could underlie previously described effects of systemic CB1R antagonists previously assumed to be associated to direct interaction with central targets.

## Materials and Methods

### Animals

Young adult (10-12 weeks old) male Swiss albino (CD-1) mice (Charles River, France) were used for pharmacological approaches in behavioural, microdialysis, electrophysiological and biochemical experiments. For genetic approaches to reduce CB1R expression, heterozygous mice for the *Cnr1* gene (CB1HZ) and their wild-type littermates in C57BL/6J genetic background were used (Zimmer et al., 1999). Conditional KO mice for the *Cnr1* gene lacking CB1R exclusively in DBH-expressing cells were generated as previously detailed (Busquets-Garcia et al., 2016) in C57BL/6J genetic background.

Mice were housed in Plexiglas cages (2-4 mice per cage) and maintained in temperature (21 ± 1 °C) and humidity (55 ± 10%) controlled environment. Food and water were available *ad libitum*. All the experiments were performed during the light phase of a 12 h cycle (light on at 8 am; light off at 8 pm). Mice were habituated to the experimental room and handled for 1 week before starting the experiments. All behavioural experiments were conducted by an observer blind to experimental conditions.

All animal procedures were conducted following “Animals in Research: Reporting Experiments” (ARRIVE) guidelines and standard ethical guidelines (Kilkenny et al., 2010) (European Directive 2010/63/EU). Animal procedures were approved by the local ethical committee (Comitè Ètic d’Experimentació Animal-Parc de Recerca Biomèdica de Barcelona, CEEA-PRBB).

### Drugs and treatments

Rimonabant (Axon Medchem) and mifepristone (Sigma-Aldrich) were dissolved in 5% ethanol, 5% cremophor-EL and 90% saline (0.9% NaCl). AM6545 (Tocris-Bio-Techne) was dissolved in 0.26% DMSO, 4.74% ethanol, 5% cremophor-EL and 90% saline. Sotalol (Sigma-Aldrich) was dissolved in saline. Propranolol (Sigma-Aldrich) was dissolved in saline. For the activation of the inhibitory designer receptors exclusively activated by designer drugs (hM4Di-DREADD), clozapine N-oxide (CNO) (Enzo Life Sciences, NY) was dissolved in sterile saline. All intraperitoneally (i.p.)-injected drugs were administered in a volume of 10 mL/kg of body weight at the doses and time points indicated.

### Viral vectors

We used the following vectors: AAV-hM4Di-DREADD (AAV5-hSyn-hM4D(Gi)-mCherry) and AAV-control-DREADD (AAV5-hSyn-mCherry) from Addgene.

To detect the viral expression in all the experiments we visualized through epifluorescence microscopy the reporter mCherry in the nodose ganglion.

### Bilateral adrenalectomy

Mice were anesthetized by isoflurane inhalation, 5% v/v induction and 3% v/v for maintenance, with oxygen (0.8 L/min). A small incision of 1 cm was made in the left and right flanks, and the adrenal glands were identified and removed from the surrounding tissue. Wounds were closed in two layers using 4/0 silk sutures (Alcon). All animals were given access to saline after surgery to ensure adequate salt balance. The experiments were resumed following a recovery period of 10 d.

### Bilateral intra-hippocampal cannula implantation

Bilateral intra-hippocampal cannula implantation was performed as previously described (Busquets-Garcia et al., 2018). Briefly, mice were deeply anaesthetized with a mixture of ketamine hydrochloride (Imalgène; Merial Laboratorios S.A.) and medetomidine hydrochloride (Domtor; Esteve, Spain) dissolved in sterile 0.9% physiological saline and administered intraperitoneally (i.p., 75 mg/kg, and 1 mg/kg of body weight respectively). Mice were placed in a stereotaxic frame (David Kopf, Tujunga, CA) and an incision was made over the skull and burr holes were drilled with the size of the guide cannula. A bilateral 26-gauge guide cannula (Plastics One, Roanoke, VA) was implanted into the dorsal hippocampus as a guide for a bilateral injection cannula (33-gauge internal cannula, Plastics One). The bilateral guide cannula was fixed using dental cement (Dentalon plus, Heraeus Kulzer GmbH, Hanau, Germany) and two stainless-steel screws. The placement was set at 1 mm above the target injection site and the guide cannula was sealed with a dummy of stainless-steel wire with 0.5 mm of projection to prevent obstruction. The target injection site coordinates were as follows: anteroposterior, −1.80 mm; mediolateral, ±1.00 mm; dorsoventral, 2.00 mm relative to bregma (Paxinos and Franklin, 2001).

After surgery, anesthesia was reversed by s.c. atipamezole administration (Antisedan, Ecuphar, Spain). Moreover, mice received i.p. administration of gentamicine and s.c. administration of meloxicam.

Animals were kept on a 37 °C heating pad during the surgery, and until recovery from anesthesia. The behavioral experiments started 1 week after surgery.

### Intra-hippocampal drug administration

After the familiarization phase of NORT, mice received a bilateral intra-hippocampal injection of 0.50 μL of propranolol (0.5, 1 or 2 μg) or saline at a constant rate of 0.25 μL/min by using a microinfusion pump during 2 min. The injection cannula projected 1.00 mm below the ventral tip of the implanted guide cannula. The displacement of an air bubble inside the length of the polyethylene tubing that connected the Hamilton syringe to the injection needle was used to monitor the microinjections. After infusion, the injection cannula was left for an additional period of 2 min to allow the fluid to diffuse and to prevent reflux before withdrawal.

### Verification of cannula position

After behavioural evaluation, mice were killed by cervical dislocation and the brains removed and frozen at -80°C. Coronal slices were obtained on a cryostat, mounted on slides and stained with cresyl violet. The injection sites were verified using a light microscope in blinding conditions. All mice with cannula located outside the hippocampus were excluded from the experiment.

### Surgery and virus vector microinjection in the vagus nerve

Mice were anesthetized with a mixture of ketamine hydrochloride (Imalgène; Merial Laboratorios S.A.) and medetomidine hydrochloride (Domtor; Esteve, Spain) dissolved in sterile physiological saline and administered intraperitoneally (i.p., 75 mg/kg, and 1 mg/kg of body weight respectively) and placed in a supine position. A small incision from the larynx to approximately the beginning of the sternum was made in the left and right flanks. Submaxillary glands were esposed by blunt dissection and the midline was opened to expose the trachea. Vagus nerve and the carotid artery were identified within the set of neurovascular fibers parallel to the trachea. The nerve was isolated from the artery using a spinal hook, taking great care not to damage the artery.

To keep the nerve fixed, a pair of blunt-tipped forceps were introduced between the nerve and the rest of the tissue. Then, 1µL of a viral suspension (AAV) was very slowly inoculated using a Hamilton syringe (34G, 45° bevel). Finally, the holding forceps were removed and the nerve was returned to its initial position. The same procedure was repeated on the nerve of the other side. Once the inoculations finished, the exposed area was closed in layers and the skin incision was closed by means of a simple stitch suture with 4/0 braided silk. Then, anesthesia was reversed by s.c. atipamezole administration (Antisedan, Ecuphar, Spain).

The animals were then administered for 48 hours with analgesic and antibiotic. A possible nerve injury was also monitored by analysis of urination and defecation capacity.

### Novel object recognition test

The Novel object recognition test (NORT) was performed following a protocol previously described (Gomis-González et al., 2021).

On the first day, mice were habituated to the V-shaped maze (V maze) for 9 min. Next day, 2 identical objects (familiar objects) were presented at the end of each corridor of the V-maze and mice were left to explore for 9 min (familiarization phase). On the last day, 24h or 48h after the familiarization phase, one of the familiar objects was replaced by a new object (novel object) and mice were placed back in the V-maze to evaluate memory performance (test phase). In this phase, the time spent exploring each of the objects (familiar and novel) was measured to calculate a discrimination index (DI = (Time Novel Object -Time Familiar Object)/(Time Novel Object + Time Familiar Object)), defining exploration as the orientation of the nose toward the object at a distance closer than 1 cm. A higher discrimination index is considered to reflect greater memory retention for the familiar object. Mice that explored <10 sec both objects during the test session or <2 sec one of the objects were excluded from the analysis. Drug treatment was administered after the familiarization phase.

### Locomotor activity

Locomotor activity was assessed for 120 min after acute administration of AM6545. Individual locomotor activity boxes (9 × 20 × 11 cm) (Imetronic) were used in a dim environment. The total activity and the total number of rearings were detected by infrared sensors to detect locomotor activity and infrared plane to detect rearings.

### Tissue preparation for immunofluorescence

Mice were deeply anesthetized by i.p. injection (0.2 mL/10 g of body weight) of a mixture of ketamine/xylazine (100 mg/kg and 20 mg/kg, respectively) prior to intracardiac perfusion of cold 4% paraformaldehyde in 0.1 M phosphate buffer, pH7.5 (PB) delivered with a peristaltic pump at 19 mL/min flow for 3 min. Brains were removed and post-fixed overnight at 4 °C in the same fixative solution. The next day, brains were moved to PB at 4 °C. Coronal brain sections of 30 μm were made on a freezing microtome (Leica SM200R) and stored in a 5% sucrose solution at 4 °C until use.

### Immunofluorescence

Free-floating brain slices were rinsed in PB and blocked in a solution containing 3 % donkey serum (DS) (Sigma-Aldrich) and 0.3 % Triton X-100 (T) in PB (DS-T-PB) at room temperature for 2 h, and incubated overnight in the same solution with the primary antibody to c-Fos (sc-7202, 1:1000, rabbit, Santa Cruz Biotechnology) at 4 °C. Twenty four hours later, slices were rinsed 3 times in PB and incubated at room temperature with the secondary antibody AlexaFluor-555 donkey anti-rabbit (1:1000, Jackson ImmunoResearch) in DS-T-PB for 2 h. Then, sections were rinsed and mounted onto gelatin-coated slides with Mowiol mounting medium.

### Image acquisition and cell quantification

Immunostained brain sections were analysed with a ×10 objective using a Leica DMR microscope (Leica Microsystems) equipped with a digital camera Leica DFC 300FX (Leica Microsystems). Seven different brain subregions were analysed: prelimbic cortex (PL), infralimbic cortex (IL), dentate gyrus (DG), Cornu Ammonis 1 (CA1) and 3 (CA3), basal amygdala (BA), central amygdala (CeA) and lateral amygdala (LA). For PL and IL analysis, a 430-μm-sided square region of interest (ROI) was delimited for quantification. For the rest of regions, the DAPI signal was used as counterstaining to identify and delimitate each area for quantification. The images were processed using the ImageJ analysis software. c-Fos-positive neurons in each brain area were quantified using the automatic “particle counting” option with a fixed configuration that solely detected c-Fos-positive cell bodies matching common criteria of size and circularity. A fixed threshold interval was set to distinguish the c-Fos-positive nuclei from the background. In addition, all quantifications were individually checked for homogeneity by an expert observer blind to the experimental conditions. 4-6 representative brain sections of each mouse were quantified, and the average number of c-Fos-positive neurons was calculated for each mouse. The data are expressed as density: the mean number of c-Fos-positive cells per squared mm (n = 6 mice per experimental group). For the c-Fos data, the displayed images were flipped for orientation consistency and transformed to inverted grey scale.

### C-Fos based network construction and graph analysis

Correlation matrices and circos plot were generated using Pearson *r* values from interregional correlations of c-fos expression data. Pearson *r* values were converted to Z-scores using Fisher’s transformation to determine effects of AM6545 on positive and negative functional connectivity and compare across experimental groups. All graph analysis was performed in R (v4.0) using the DescTools (v99.40; Andri Signorell et mult. al., 2021) and the ggcorrplot (v0.1.3; Alboukadel Kassambara, 2019) packages.

### rsfMR1: Image acquisition

Magnetic resonance image was performed in the Magnetic Resonance Core Facility (IDIBAPS 7T MRI Unit, Barcelona, Spain). Experiments were conducted on a 7.0T BioSpec 70/30 horizontal animal scanner (Bruker BioSpin, Ettlingen, Germany), equipped with an actively shielded gradient system (400 mT/m, 12 cm inner diameter). The receiver coil is a surface coil for the mouse brain. Animals were sedated (induction dose of 4% isoflurane in a mixture of 30% O_2_ and 70% N_2_) and placed in supine position in a Plexiglas holder with a nose cone for administering anesthetic gases and were fixed using tooth and ear bars and adhesive tape. Eyes were protected from dryness with Siccafluid 2.5 mg/g ophthalmologic fluid. Once placed in the holder and keeping isoflurane to 1.5 %, a subcutaneous bolus (0.3 mg/kg) of medetomidine (Domtor, Orion Pharma, Spain) was injected. For the next 15 min, the isoflurane dose was progressively decreased until 0.5 %. Then, a continuous perfusion of 0.6 mg/kg/h medetomidine started and was maintained until the end of the acquisition session. After completion of the imaging session, 1 µL/g of atipamezol per mouse weight (Antisedan, Orion Pharma, Spain) and saline were injected to reverse the sedative effect and compensate fluid loss. Localizer scans were used to ensure accurate position of the head at the isocenter of the magnet. Anatomical T2 RARE images were acquired in coronal orientation with effective TE=33 ms TR=2.3 s, RARE factor=8, voxel size = 0.08 × 0.08 mm^2^ and slice thickness = 0.5 mm. rs-fMRI was acquired with an EPI sequence with TR=2 s, TE=19.4, voxel size 0.21×0.21 mm^2^ and slice thickness 0.5 mm. 420 volumes were acquired resulting in an acquisition time of 14 min.

### rsfMR1: Image analysis and processing

Two approaches were used to evaluate functional connectivity. On the one hand, whole-brain connectivity was evaluated using global and regional network metrics (Rubinov & Sporns, 2010) and on the other hand seed-based analysis was performed to evaluate connectivity of posterior brainstem with the rest of the brain. For both approaches, rs-fMRI was preprocessed, including slice timing, motion correction by spatial realignment using SPM8, correction of EPI distortion by elastic registration to the T2-weighted volume using ANTs (Avants et al., 2008), detrend, smoothing with a full-width half maximum (FWHM) of 0.6mm, frequency filtering of the time series between 0.01 and 0.1 Hz and regression by motion parameters. All these steps were performed using NiTime (http://nipy.org/nitime). Brain parcellation was performed by registration of a mouse brain atlas (Ma et al., 2008) to the T2-RARE acquisition of each subject using ANTs diffeomorphic registration. Region parcellation was then registered from the T2-weighted volume to the preprocessed mean rs-fMRI. Whole-brain functional brain network was estimated considering the gray matter regions obtained by parcellation as the network nodes. Connectivity between each pair of regions was estimated as the Fisher-z transform of the correlation between average time series in each region. Network organization was quantified using regional and global graph metrics. Strength, local efficiency and clustering coefficient was computed for every region and whole brain connectivity was quantified by average strength, global and local efficiency and average clustering coefficient (Rubinov & Sporns, 2010). To perform the seed-based analysis, posterior brainstem containing the nuclei of the solitary tract was manually drawn in the images of each animal. Average time series in the seed was computed and correlated with each voxel time series, resulting in a correlation map describing the connectivity of the region of interest with the rest of the brain. Voxel-wise differences between the correlation maps were estimated using fsl-randomize, that performs non-parametric permutation test including family-wise error correction on neuroimaging data (Winkler et al., 2014).

### In vivo microdialysis

Animals were anesthetized with isoflurane (1.5 - 2.5 % v/v for induction and maintenance) and placed in a Kopf stereotaxic frame. Intracerebral probe (cuprophan membrane of 2 mm) was implanted in the hippocampus and fixed to the skull. The target injection site coordinates were as follows: anteroposterior, −3.40 mm; mediolateral, ±2.60 mm; dorsoventral, -4.20 mm relative to bregma (Paxinos and Franklin, 2001). The next day, mice were placed in a plastic bowl and connected to a fraction collection system for freely-moving animals (Raturn, BASi, USA). The input tube of the dialysis probe was connected to a syringe pump (BeeHive and BabyBee, BASi), which delivered a modified cerebrospinal fluid (CSF) containing NaCl 148 mM, KCl 2.7 mM, CaCl2 1.2mM and MgCl2 0.85 mM (pH 7.4) to the probe at a rate of 1 µL/min. The output tubes from the animals were attached to a refrigerated fraction collector (HoneyComb, BASi). Samples were collected every 35 min for the analysis of the different neurotransmitters in vials containing 5 µL of acetic acid 0.1 M. Eight baseline samples were collected from each animal, but only the last 6 ones were used for subsequent analysis.

### NA, DA and 5-HT chromatographic analysis

Neurotransmitter concentrations were measured immediately after samples collection by Ultra Performance Liquid Chromatography (UPLC) coupled to an electrochemical detector (Alexys analyser, Antec Leyden, Holland). The mobile phase consisted of 100 mM phosphoric acid, 100 mM citric acid, 0.1 mM EDTA, 950-1500 mg/L 1-octanesulfonic acid (OSA), 5% v/v acetonitrile; the pH was adjusted to 6.0 with 50% NaOH/ 45% KOH solution. The flow rate of the mobile phase was 0.075 mL/min and the temperature for the analytical column (Acquity UPLC BEH C18, 1.7 µm, 1×100 mm; Waters, Milford, USA) was 37 °C.

### In vivo electrophysiological recording

Mice were anaesthetized with uretano (1.2 g/kg, i.p.), and placed in the stereotaxic frame with the skull positioned horizontally. A burr hole was drilled and the recording site coordinates were as follows: anteroposterior, -1.5 mm; mediolateral, ±0.2-1.2 mm; dorsoventral, -2.7-4.0 mm relative to lambda (Gobbi et al., 2007). The body temperature was maintained at 37 °C for the entire experiment using a heating pad.

Single-unit extracellular recordings of mouse LC neurons were performed as previously described (Gobbi et al., 2007). The recording electrode was filled with 2% solution of Pontamine Sky Blue in 0.5% sodium acetate and broken back to a tip diameter of 1–2 mm. The electrode was lowered into the brain by means of a hydraulic microdrive (model 640; David Kopf Instruments). LC neurons were identified by standard criteria, which included spontaneous activity displaying a regular rhythm and firing rate between 0.5 and 5 Hz, characteristic spikes with a long-lasting (>2 ms), positive–negative waveform action potentials and the biphasic excitation–inhibition response to pressure applied on contralateral hind paw (paw pinch), as previously described in mice (Gobbi et al., 2007) and rats (Cedarbaum & Aghajanian, 1978). The extracellular signal from the electrode was pre-amplified and amplified later with a high-input impedance amplifier and then monitored on an oscilloscope and on an audio monitor. This activity was processed using computer software (Spike2 software; Cambridge Electronic Design) and firing rate was calculated. Basal firing rate and other electrophysiological parameters were measured for 3 min. Changes in firing rate were expressed as percentages of the basal firing rate (mean firing rate for 3 min prior to drug injection) and were measured after 35 min until the end of the experiment. Only 1 cell was studied in each animal when any drug was administered.

### Statistical analysis

Results are reported as mean ± standard error of the mean (s.e.m). Data analysis were performed using GraphPad Prism software (GraphPad Software). Statistical comparisons were evaluated using unpaired Student’s t-test for 2 groups comparisons or two-way ANOVA for multiple comparisons. Subsequent Tukey *post hoc* was used when required (significant interaction between factors). Comparisons were considered statistically significant when *p* < 0.05.

## Acknowledgements

We thank Dulce Real, Marta Linares and Francisco Porrón for expert technical assistance, and the Laboratory of Neuropharmacology-NeuroPhar for helpful discussion. S.M-T. was the recipient of a predoctoral fellowship (Generalitat de Catalunya) [FI-B00531 2016], A.B-M. was supported by predoctoral fellowship (Generalitat de Catalunya) [FI_B0052 2020]. This study was supported by Ministerio de Economía, Innovación y Competitividad (MINECO), Spain (#RTI2018-099282-B-I00B to A.O., # PID2020-120029GB-I00 to R.M.; Generalitat de Catalunya, Spain (2017SGR-669 to R.M.); Basque Government, Spain (IT-1211-19 to J.J.M); ICREA (Institució Catalana de Recerca i Estudis Avançats, Spain) Academia to A.O. and R.M. Grant “Unidad de Excelencia María de Maeztu”, funded by the MINECO (#MDM-2014-0370). FEDER, European Commission funding is also acknowledged.

This research was supported by PRPSEM Project with ref. RTI2018-099773-B-I00 from (MCIU/FEDER/AEI), the CERCA Program, and the Commission for Universities and Research of the Department of Innovation, Universities, and Enterprise of the Generalitat de Catalunya (SGR2017-648) to JADR. IBEC is the recipients of a Severo Ochoa Award of Excellence from MINECO (CEX2018-000789-S). SMT and AH were supported by Severo Ochoa program at IBEC.

S.M-T. and A.B-M. performed the behavioral, biochemical, histological, confocal and surgery experiments, statistical analyses and graphs and wrote the manuscript. L.G-L. performed bilateral hippocampal cannula implantation in mice. F.R. and B.L. generated and provided the conditional transgenic mice (DBH-CB1KO). A.O-A. performed adrenalectomy in mice. A.H. and JA.dR. designed and performed the chemogenetic experiment. A.H. performed chemogenetic surgeries in the vagus nerve. E.M-M. and G.S. performed and analyzed the rsfMRI experiments and conectomics on c-Fos-based network. JE.O. and JJ.M. performed the microdyalisis in vivo experiments in mice. JA.R-O. designed and performed in vivo recordings in mice. R.M. supervised the study; A.O. conceptualized, supervised the study and wrote the manuscript. All authors revised the final version of the manuscript.

---

**Supplementary Fig. 1.**
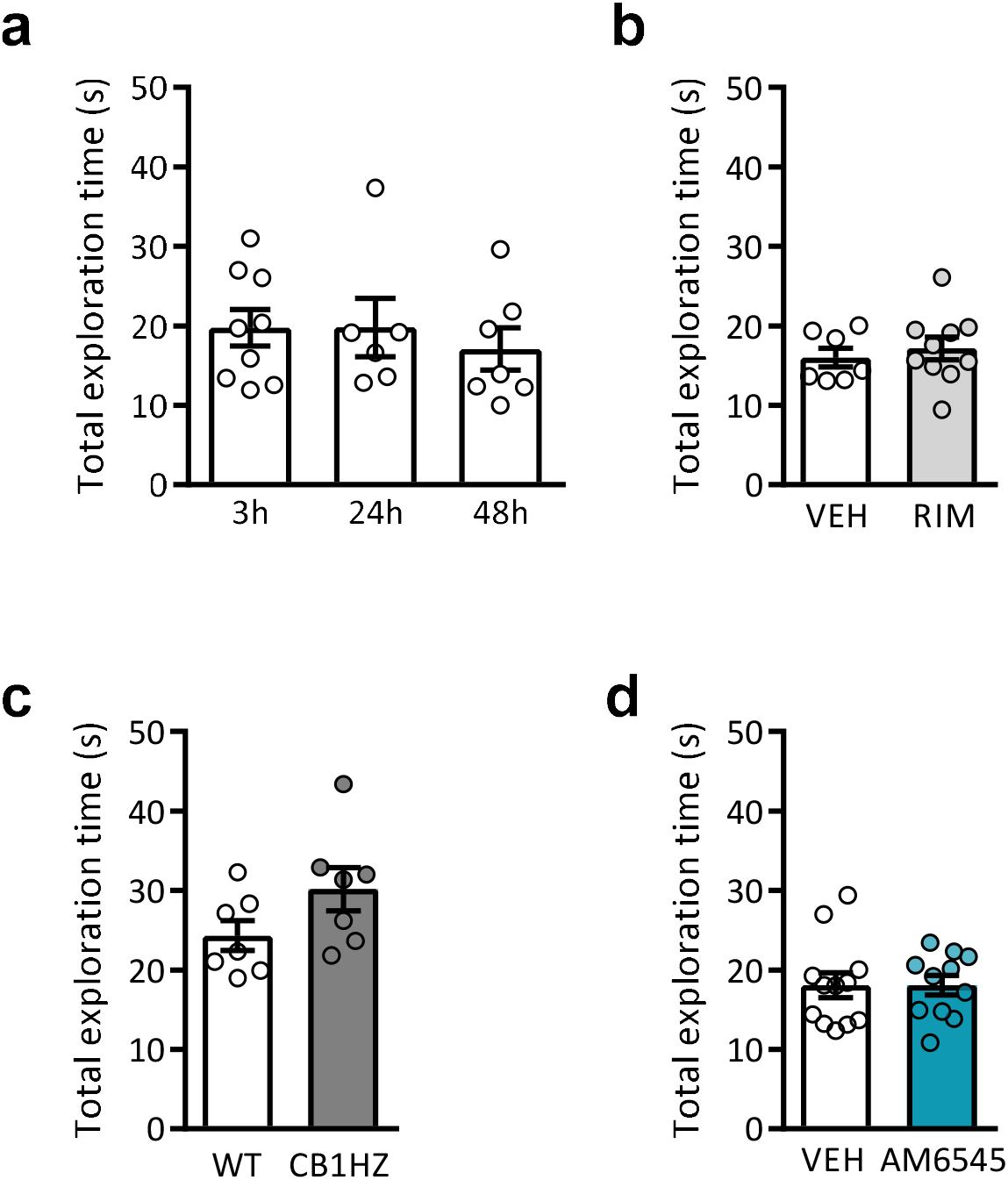
Total exploration times in NORT after pharmacological or genetic CB1R inhibition. **a** Total exploration times obtained at 3 h, 24 h and 48 h after the familiarization phase (n = 5 - 8) **b-d** Total exploration times in NORT at 48h **b** after acute post-familiarization treatment with vehicle (VEH) or rimonabant (RIM) (1 mg/kg) (n = 7 - 11) **c** in CB1HZ and WT mice (n = 6 - 8) **d** after acute post-familiarization treatment with vehicle (VEH) or AM6545 (1 mg/kg) (n = 7 - 8). Data are expressed as mean ± s.e.m. Statistical significance was calculated by one-way ANOVA test or Student’s t test.

**Supplementary Fig. 2.**
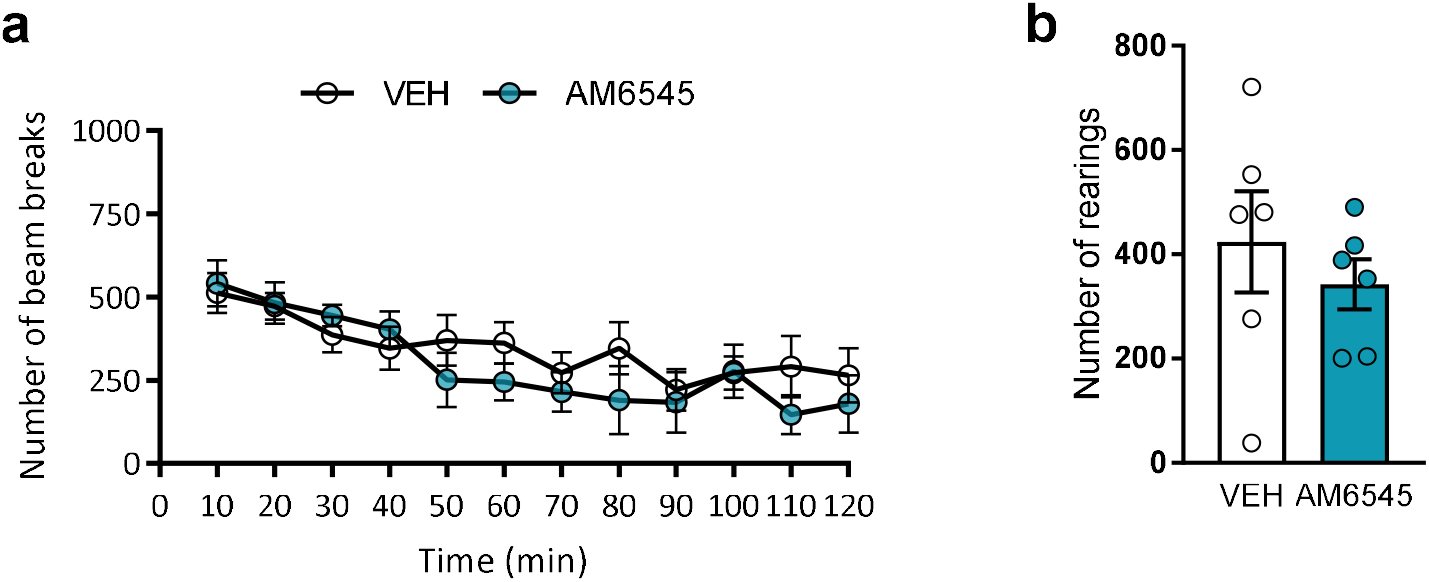
Locomotor activity after acute AM6545 administration. **a** Total activity and **b** total number of rerarings performed in locomotor activity boxes for 120 min by mice treated with vehicle (VEH) or AM6545 (1 mg/kg) (n = 6). Data are expressed as mean ± s.e.m. Statistical significance was calculated by two-way repeated measures ANOVA test or Student’s t test.

**Supplementary Fig. 3.**
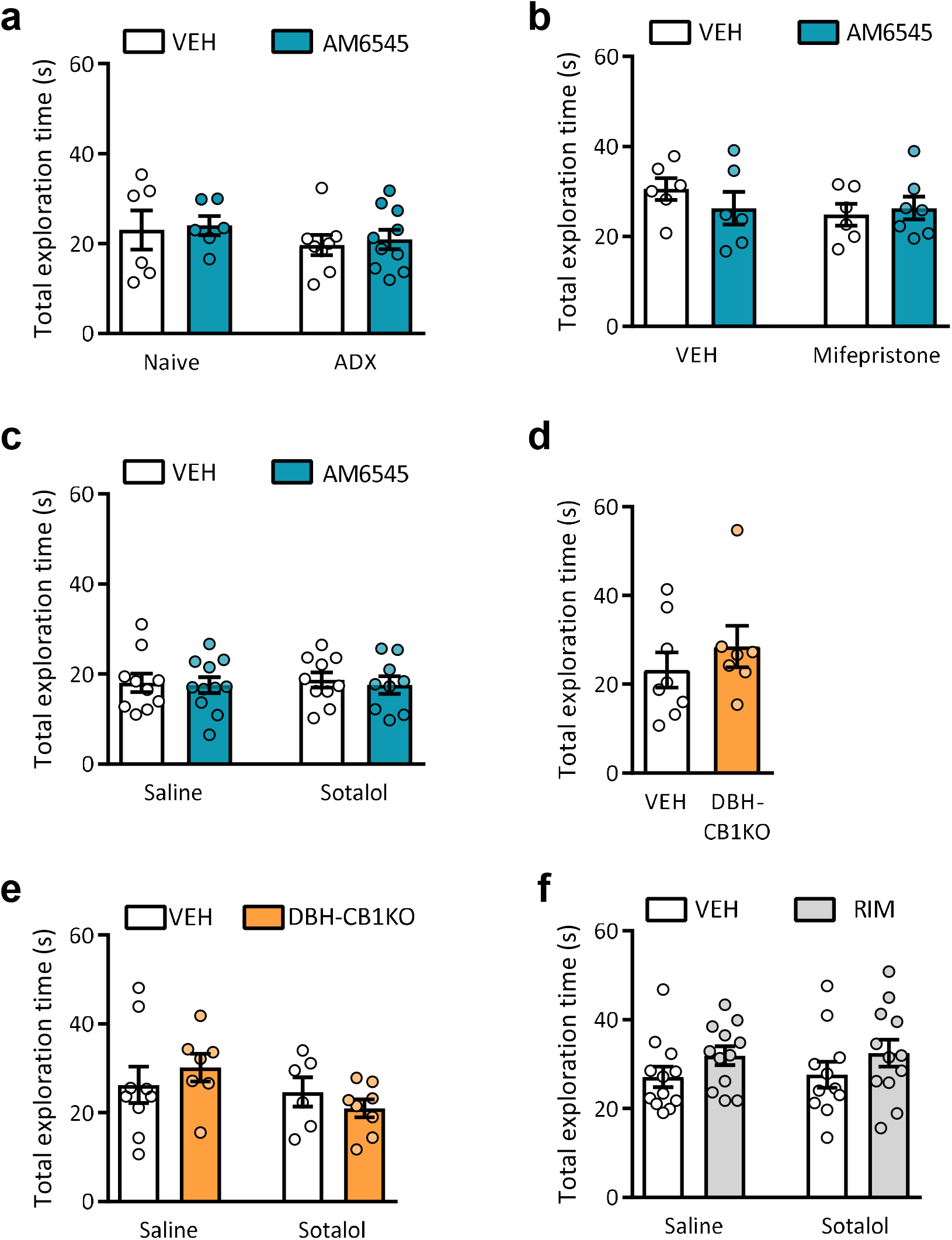
Total exploration times of systemic or peripheral CB1R blockade are not affected. **a-f** Total exploration times obtained in the NORT performed at 48 h of **a** adrenalectomized (ADX) or naive mice treated with vehicle (VEH) or AM6545 (1 mg/kg) (n = 6 - 8), **b**,**c** mice treated with vehicle (VEH) or AM6545 (1 mg/kg) after pre-treatment with **b** vehicle (VEH) or mifepristone (50 mg/kg) (n = 6 - 7) and **c** saline or sotalol (10 mg/kg) (n = 8 - 10), **d** WT or DBH-CB1KO mice (n = 5 - 6), **e** WT or DBH-CB1KO mice after saline or sotalol (10 mg/kg) treatment (n = 6 - 8), **f** mice pre-treated with saline or sotalol (10 mg/kg) prior to rimonabant (RIM) (1 mg/kg) or vehicle (VEH) (n = 9 -11). Data are expressed as mean ± s.e.m. Statistical significance was calculated by two-way ANOVA test or Student’s t test.

**Supplementary Fig. 4.**
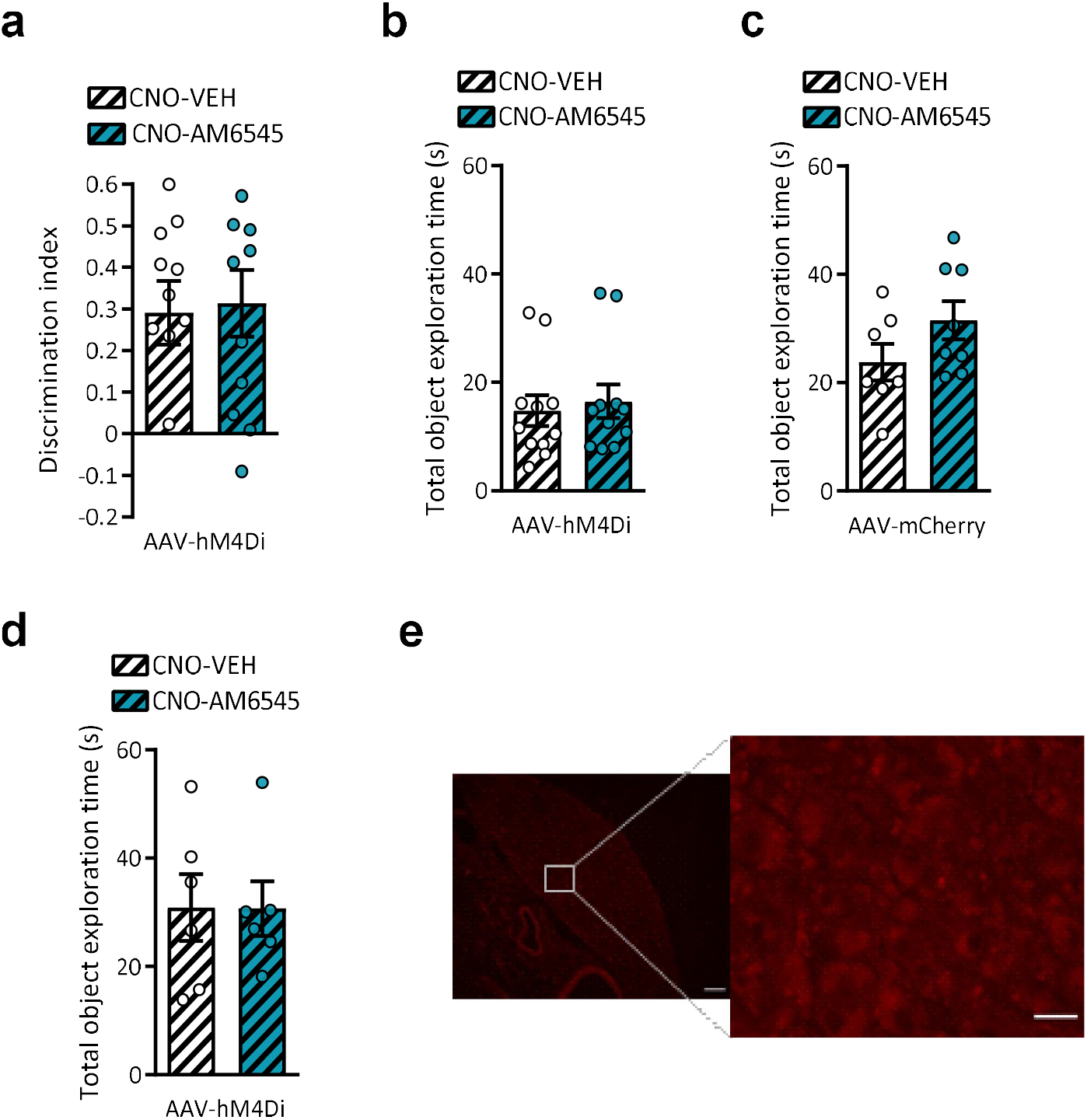
Chemogenetic targeting of the vagus nerve does not modify the behavioral responses in the NORT. **a** Discrimination index values and **b** total exploration times in NORT at 24 h after the familiarization phase and after CNO (3 mg/kg) and vehicle (VEH) or AM6545 (1 mg/kg) in mice infected with AAV-hm4Di in the vagus nerve and i.p. CNO (3 mg/kg) administration (n = 10 - 11) **c**,**d** Total exploration times in NORT at 48 h after CNO (3 mg/kg) and vehicle (VEH) or AM6545 (1 mg/kg) administration in mice infected with **c** AAV-mCherry (n = 6 - 8) or **d** AAV-hM4Di (n = 6 - 8). **e** Representative images of mCherry expression detected in nodose ganglion of mice after AAV5-mCherry injection in the vagus nerve (left image scale bar = 100µm, right image scale bar = 30µm). Data are expressed as mean ± s.e.m. Statistical significance was calculated by Student’s t test.

**Supplementary Fig. 5.**
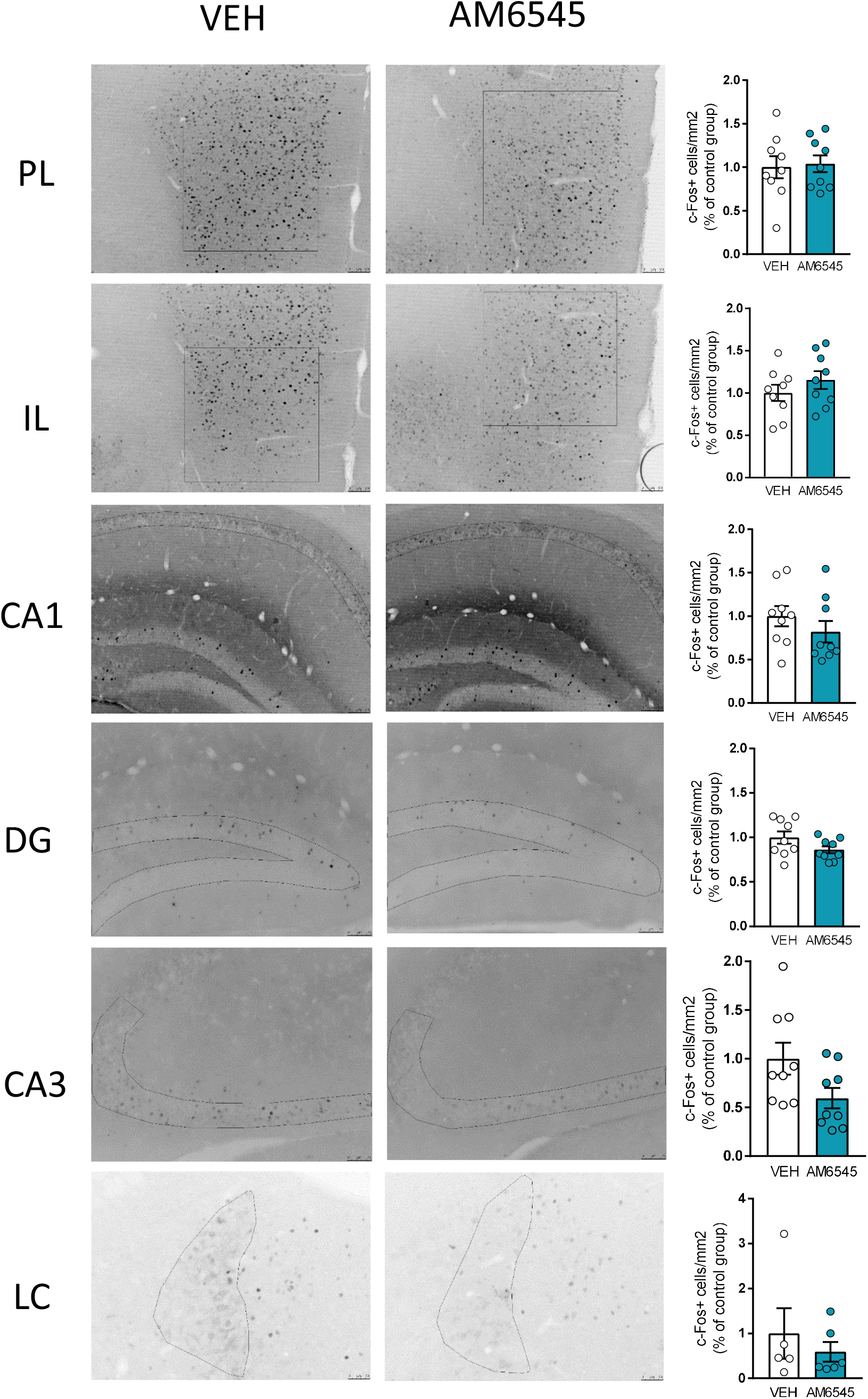
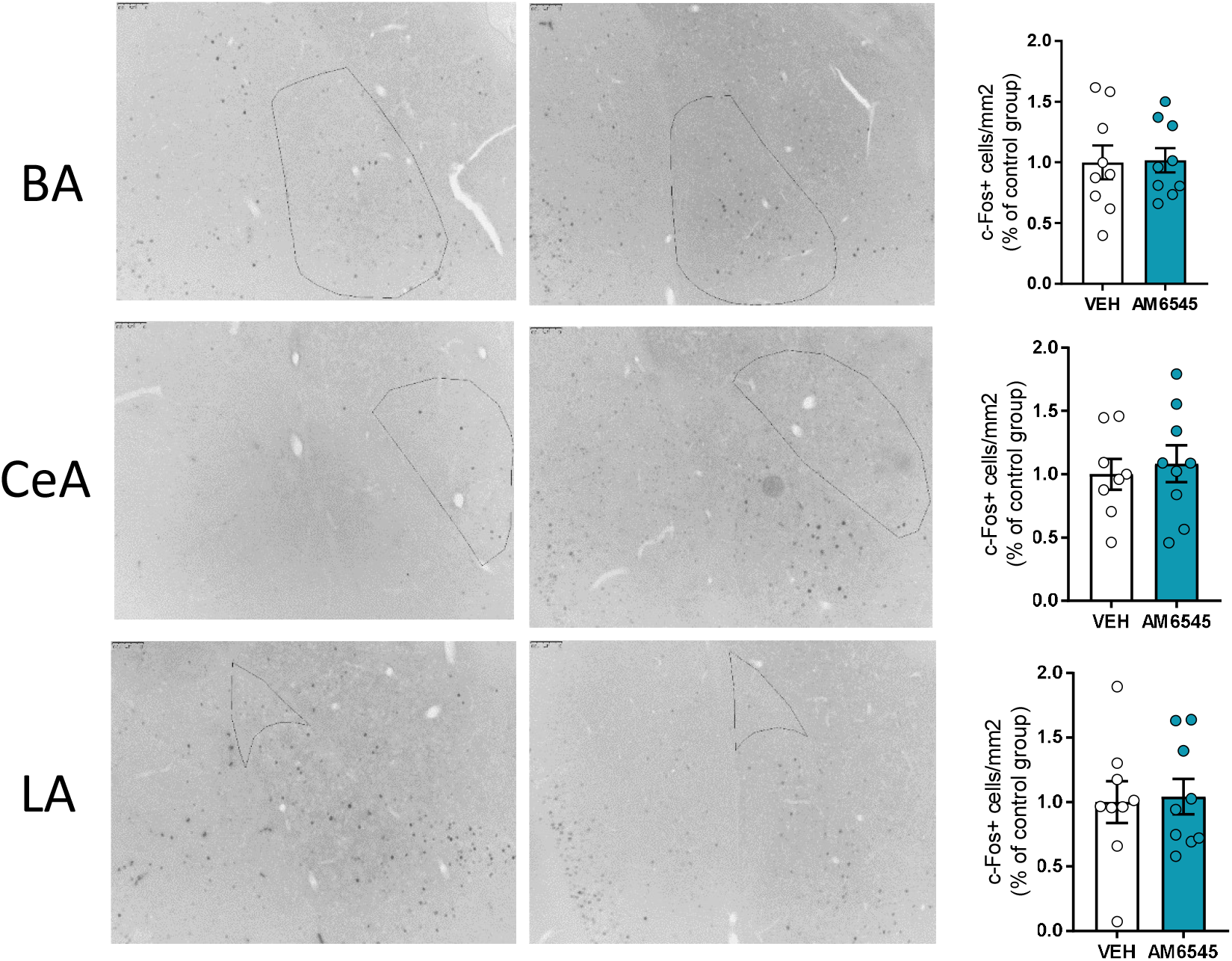
Effects of acute AM6545 treatment in c-Fos expression in specific regions of the CNS. Representative immunohistochemical staining of c-Fos positive cells 90 min after acute vehicle (VEH) or AM6545 (1 mg/kg) administration in mice (n = 5-9). Brain regions evaluated: prelimbic cortex (PL), infralimbic cortex (IL), *cornu ammonis* 1 (CA1), dentate gyrus (DG), *cornu ammonis* 3 (CA3), *locus coeruleus* (LC), basal amygdala (BA), central amygdala (CeA) and lateral amygdala (LA). Data are expressed as mean ± s.e.m. Statistical significance was calculated by Student’s t test.

**Supplementary Fig. 6.**
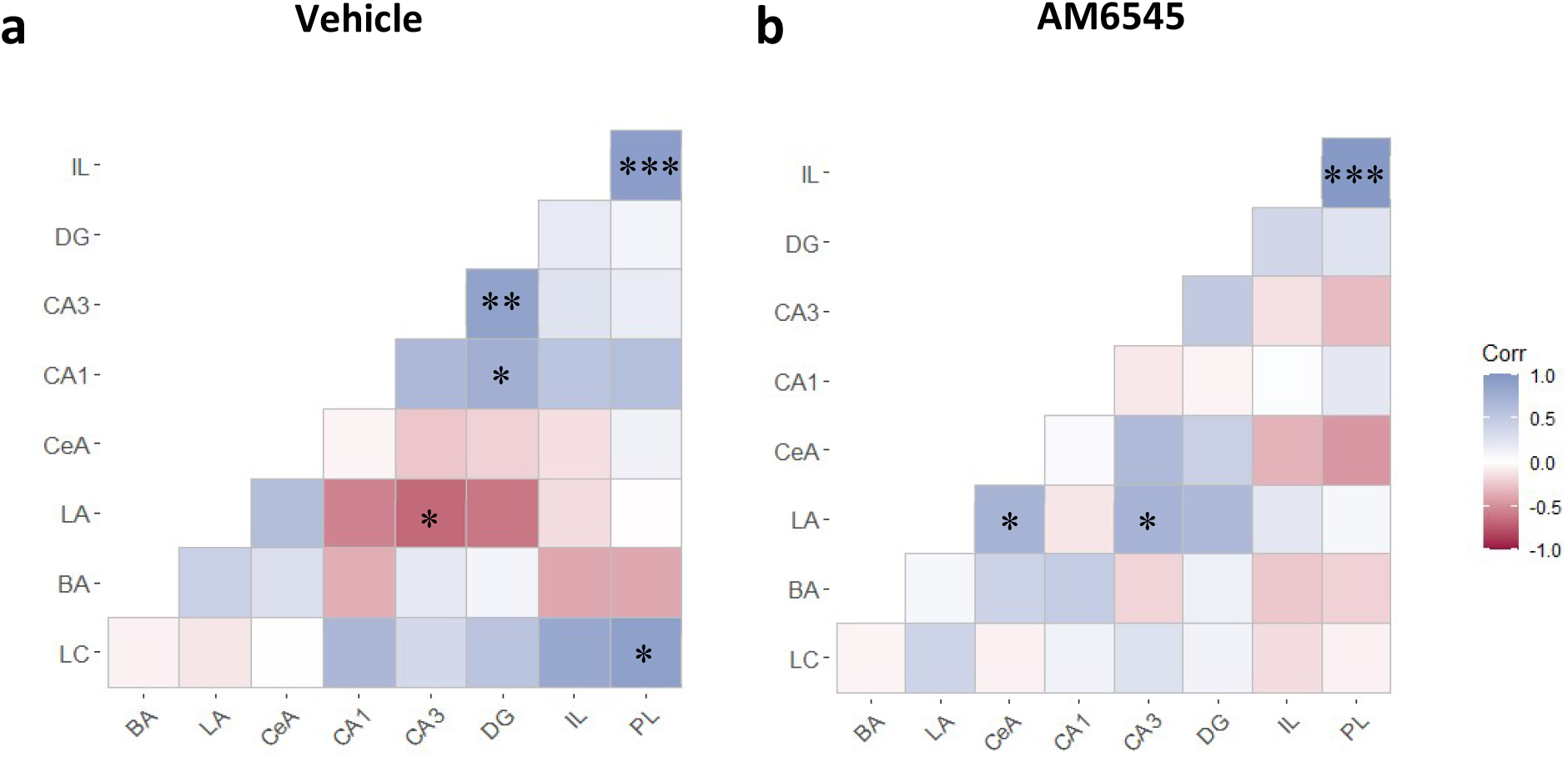
Correlation heat map of c-Fos expression in different brain regions after acute AM6545 administration. Pearson correlation analysis of c-Fos expression in **a** vehicle and **b** AM6545 (1 mg/kg) between different brain areas, including prelimbic cortex (PL), infralimbic cortex (IL), *cornu ammonis* 1 (CA1), dentate gyrus (DG), *cornu ammonis* 3 (CA3), *locus coeruleus* (LC), basal amygdala (BA), central amygdala (CeA) and lateral amygdala (LA). *p < 0.05, **p < 0.01, ***p < 0.001.

**Supplementary Fig. 7.**
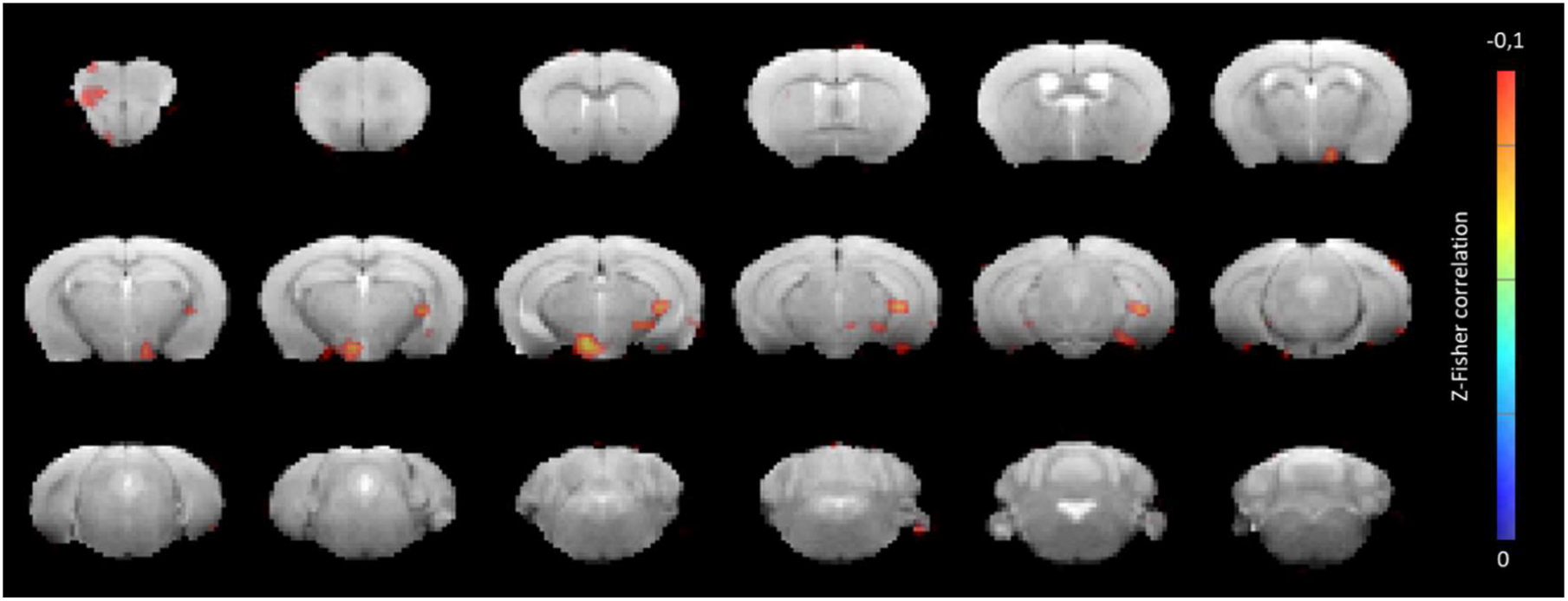
Brainstem connectivity map in AM6545-treated mice: only areas where BOLD signal is negatively correlated to the brainstem signal are shown. Color represents z-Fisher of the correlation coefficient.

**Supplementary Fig. 8.**
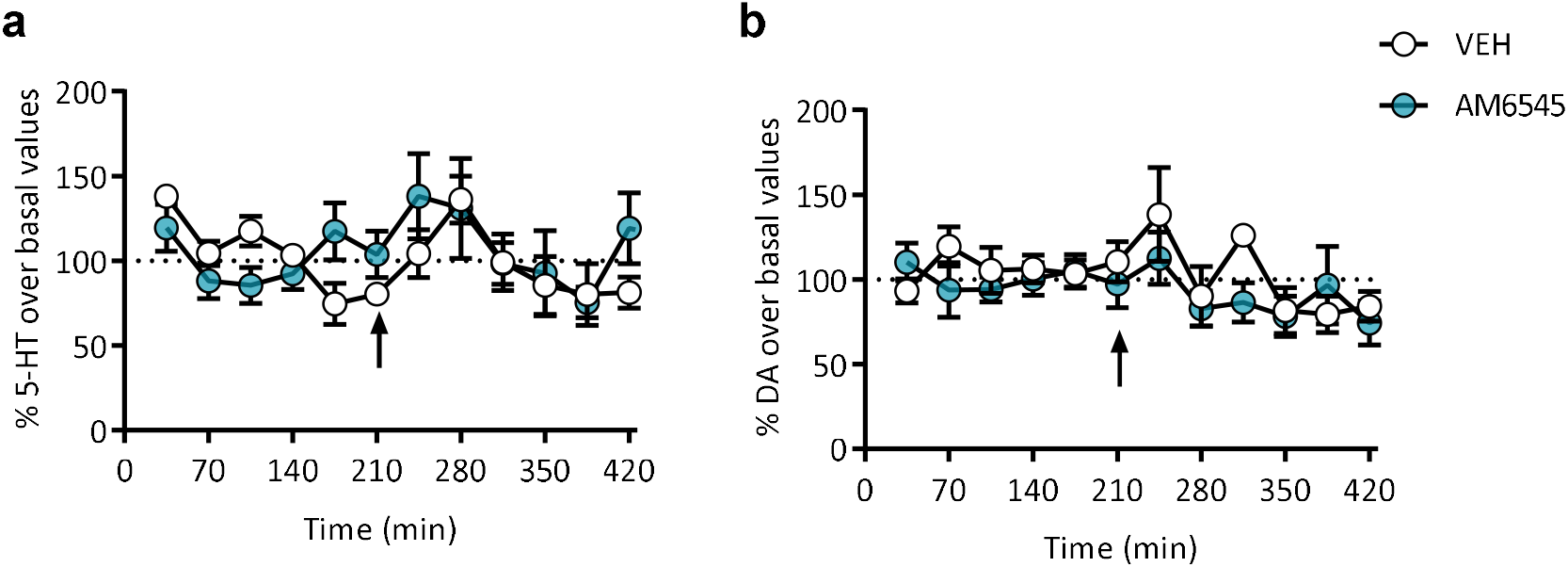
Acute AM6545 administration does not modify other monoamine extracellular levels in the hippocampus. **a-b** Extracellular **a** serotonin (5-HT) and **b** dopamine (DA) levels in the hippocampus after acute AM6545 (1 mg/kg) or vehicle (VEH) administration relative to baseline values (n = 6 - 7). Arrow indicates the time of administration. Points represent mean ± s.e.m and are expressed as percentages of baseline. Statistical significance was calculated by two-way repeated measures ANOVA test.

**Supplementary Fig. 9.**
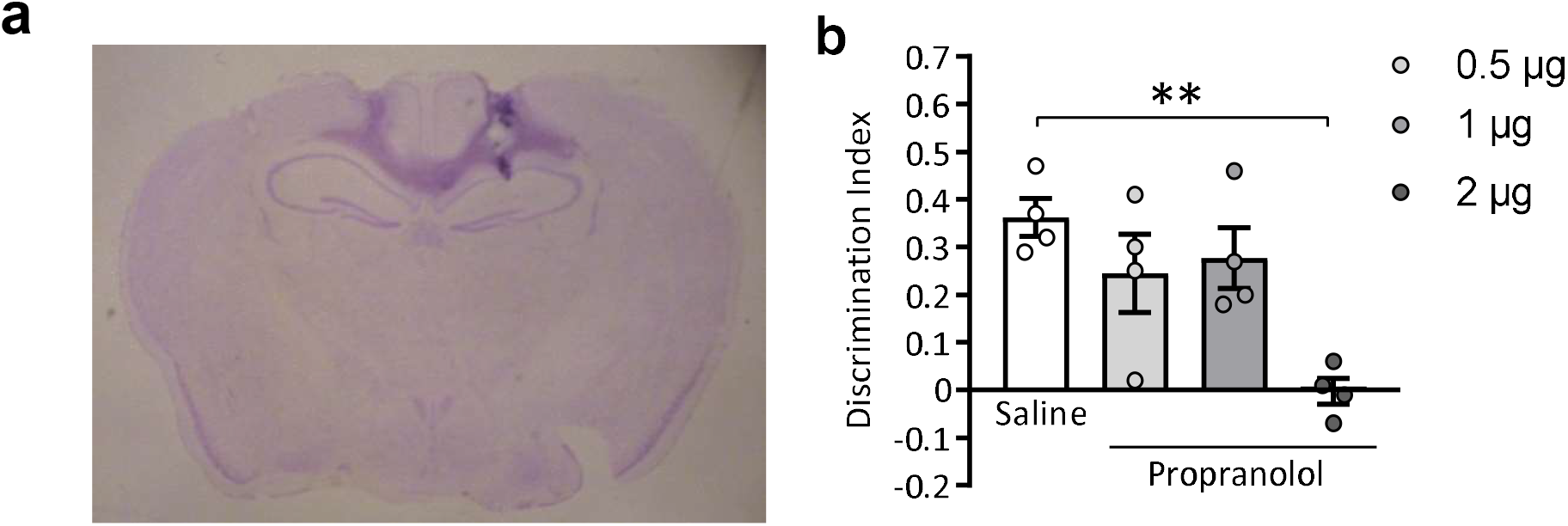
Intrahippocampal propranolol injection. **a** Cannula placement in the dorsal hippocampus. A brain coronal section from a representative mouse showing cannula placement in the dorsal hippocampus. Brain slices were stained with cresyl violet. **b** Discrimination index values of different doses of intrahippocampal propranolol infusion or saline in the NORT at 24 h (n = 4). Data are expressed as mean ± s.e.m. **p < 0.01 by one-way ANOVA test followed by Tukey *post hoc*.

